# Multiple hybridization events punctuate the evolutionary trajectory of *Malassezia furfur*

**DOI:** 10.1101/2021.11.02.466935

**Authors:** Bart Theelen, Verónica Mixão, Giuseppe Ianiri, Joleen Goh Pei Zhen, Jan Dijksterhuis, Joseph Heitman, Thomas L. Dawson, Toni Gabaldón, Teun Boekhout

## Abstract

*Malassezia* species are important fungal skin commensals and are part of the normal microbiota of humans and other animals. However, under certain circumstances these fungi can also display a pathogenic behaviour. For example, *Malassezia furfur* is a common commensal of human skin, and yet is often responsible for skin disorders but also systemic infections. Comparative genomics analysis of *M. furfur* revealed that some isolates have a hybrid origin, similar to several other recently described hybrid fungal pathogens. Because hybrid species exhibit genomic plasticity that can impact phenotypes, we sought to elucidate the genomic evolution and phenotypic characteristics of *M. furfur* hybrids in comparison to their parental lineages. To this end, we performed a comparative genomics analysis between hybrid strains and their presumptive parental lineages, and assessed phenotypic characteristics. Our results provide evidence that at least two distinct hybridization events occurred between the same parental lineages, and that the parental strains may have originally been hybrids themselves. Analysis of the mating-type locus reveals that *M. furfur* has a pseudobipolar mating system, and provides evidence that after sexual liaisons of mating compatible cells, hybridization involved cell-cell fusion leading to a diploid/aneuploid state. This study provides new insights into the evolutionary trajectory of *M. furfur* and contributes with valuable genomic resources for future pathogenicity studies.

**Importance:** *Malassezia furfur* is a common commensal member of human/animal microbiota that is also associated with several pathogenic states. Recent studies report involvement of *Malassezia* species in Crohn’s disease, a type of inflammatory bowel disease, pancreatic cancer progression, and exacerbation of Cystic fibrosis. A recent genomics analysis of *M*. *furfur* revealed the existence of hybrid isolates and identified their putative parental lineages. In this study, we explored the genomic and phenotypic features of these hybrids in comparison to their putative parental lineages. Our results revealed the existence of a pseudobipolar mating system in this species and showed evidence for the occurrence of multiple hybridization events in the evolutionary trajectory of *M. furfur*. These findings significantly advance our understanding of the evolution of this commensal microbe and are relevant for future studies exploring the role of hybridization in the adaptation to new niches or environments, including the emergence of pathogenicity.

## Introduction

*Malassezia* are basidiomycetous yeasts that are the dominant fungal component of the healthy human skin microbiome (1, 2). Nevertheless, they can also take on pathogenic roles in various skin disorders, and have been implicated in cases of infection associated with several comorbidities (3–12). Non-culture based sequencing methods revealed *Malassezia* presence in other ecological niches, including insects, nematodes, corals, sponges, deep-sea environments, and soils (13–15), suggesting that *Malassezia* are ecologically diverse. Interestingly, *Malassezia* are evolutionarily related to fungal plant pathogens, potentially pointing towards a host shift from plants to animals, facilitated by the loss of genes coding for proteins involved in the degradation of complex carbohydrates, and expansion of lipid hydrolases required to break down lipids available on animal skin (16, 17). In *Basidiomycetes*, species with either a tetrapolar or bipolar mating system have been described (18, 19). In a tetrapolar mating system, the pheromone and pheromone-receptor locus (P/R) and the homeodomain locus (HD) are unlinked, whereas in a bipolar mating system both loci are linked and contiguous on the same chromosome which precludes recombination. An intermediate -pseudobipolar-mating system was described in three *Malassezia* species, namely *Malassezia globosa, Malassezia sympodialis*, and *Malassezia yamatoensis*. In this configuration, the P/R and HD mating-type loci are linked on the same chromosome but far enough apart from each other that recombination can still occur (16, 20, 21). The presence of mating and meiotic genes suggests that *Malassezia* are potentially capable of sexual reproduction, but mating remains hitherto unobserved, and the genus is known to propagate asexually through unipolar budding (16, 21, 22).

Our study focuses on *Malassezia furfur*, a species isolated from a wide variety of hosts, from humans (with healthy skin, skin disorders, or bloodstream infections), to a range of domestic and zoo animals (12, 15, 23–26). This species presents variable cell sizes and shapes and different ploidies and karyotypes (27–29). Indeed, previous studies reported two different karyotypes -one displaying more chromosomes (10-11 vs. 7-8) and a larger genome size (∼14 Mbp vs. 8.5 Mbp) than the other- and a high degree of genetic variation that uncovered a possible hybrid genotype (28, 30, 31). These findings were later corroborated in a genomic analysis of members of the *Malassezia* genus. This analysis revealed four *M. furfur* strains (CBS1878, CBS4172, CBS7019, CBS7710) with double genome size due to gene duplication that possibly originated from a hybridization event between members of the lineages of strains CBS7982 and CBS14141 (syn. JPLK23) (16).

The coexistence of genomic material of two diverged lineages in a single cell is generally expected to have lower fitness (32, 33). Nevertheless, several genomic mechanisms, such as recombination, often lead to loss of heterozygosity (LOH), thus erasing some of the genomic incompatibilities (34). In such a scenario, organisms carrying highly plastic genomes and unique phenotypes may survive leading to new lineages able to thrive in new environments (34–36). For instance, findings of multiple pathogenic hybrids, such as *Candida albicans* or the *Cryptococcus gattii* / *Cryptococcus neoformans* species complex, has led to the hypothesis that hybridization plays an important role in the emergence of pathogenicity (30, 37–44).

Taking advantage of the existence of known candidate parental haploid lineages, this study explored genomic and some general phenotypic features of *M. furfur* hybrids to assess the genomic aftermath of hybridization in this species and determine the existence of genomic and phenotypic alterations in these hybrids when compared to their parentals. This study reveals that *M. furfur* has a pseudobipolar mating system and shows evidence that multiple hybridization events punctuated the evolution of the species.

## Results and Discussion

### *M. furfur* AFLP patterns suggest the existence of two hybrid lineages

Previous AFLP and comparative genomics analyses suggested the existence of *M*. *furfur* hybrid strains (16, 31). In this study, we further explored the genomic patterns of these hybrids and of 17 additional strains presently classified as *M*. *furfur* (Table 1) to assess how widespread hybridization is within this species. Following the original observations of a hybrid genotype (31), we performed AFLP with two different adaptor and primer set combinations for 22 *M. furfur* strains, each combination representing different polymorphisms in restriction sites broadly spread over the genomes, giving an indication of genetic relatedness and shared or discriminating fragment sizes (Materials and Methods for more details). These AFLP analyses confirmed that the banding patterns of three previously identified hybrids (CBS1878, CBS4172, CBS7019) are indeed a combination of those of the proposed parental lineages (Figure 1) (16). Nevertheless, the hybrid strain CBS7019 did not cluster with the other two, suggesting the existence of two putative hybrid clades (Figure 1). In total, the AFLP clustering analysis identified four lineages: P1 (parental 1), which corresponds to the parental CBS7982 lineage with five additional strains; P2, which corresponds to the parental CBS14141 lineage and four additional strains; H1 (hybrid 1), which corresponds to CBS1878, CBS4172, and five additional hybrid strains; and finally, H2, which harbors CBS7019 and three additional hybrid strains (Figure 1).

**Figure 1.**
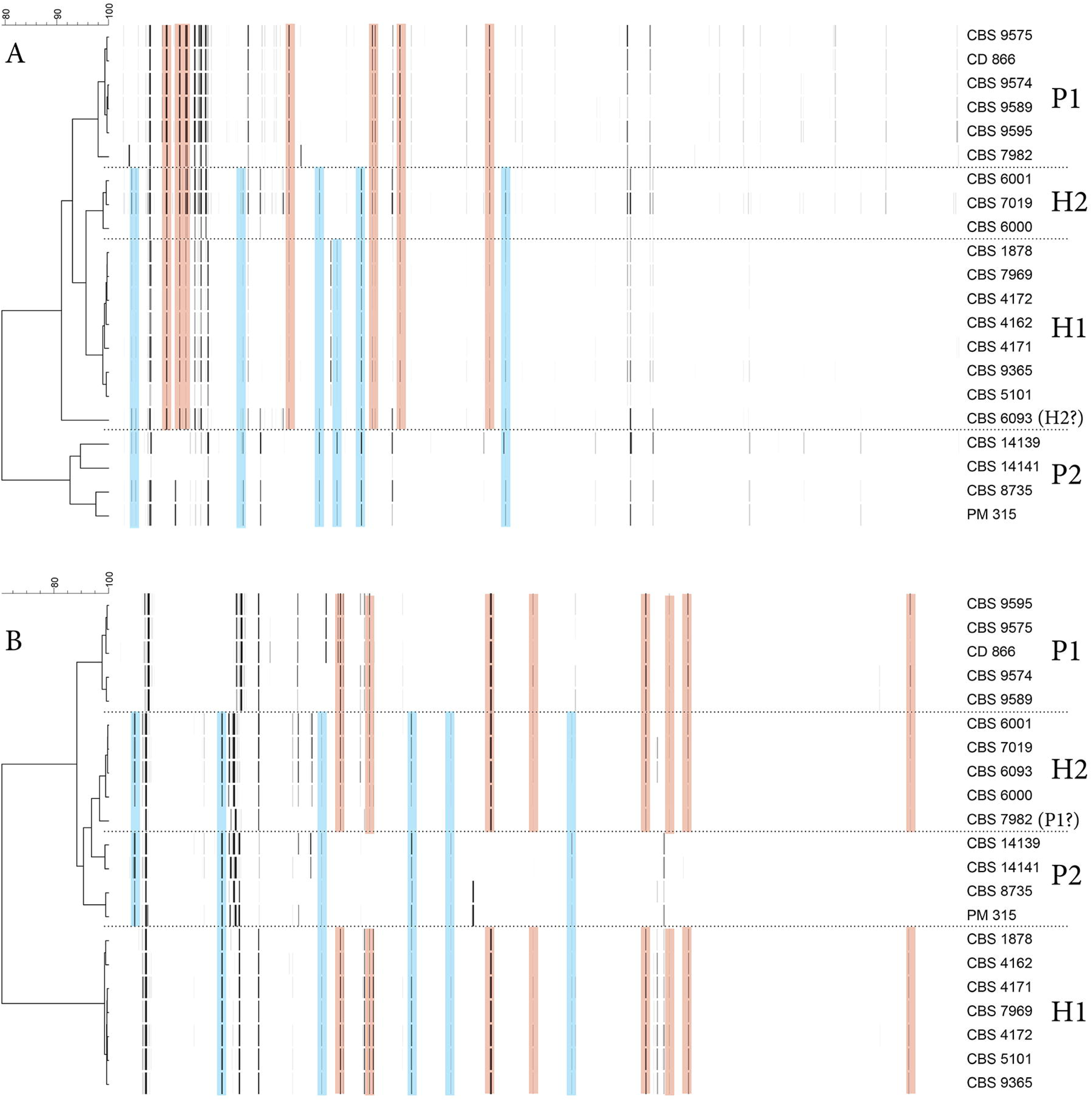
AFLP banding pattern representations derived from electropherograms with Neighbor Joining trees for two different adaptor/primer combinations (A-B) (see Materials & Methods). The horizontal scale represents the similarity percentage. Pink shading highlights restriction fragments shared between parental 1 lineage and hybrids; blue shading highlights shared restriction fragments between parental lineage 2 and hybrids. Both versions resulted from using different primer/adaptor pairs, reflecting different polymorphisms in the genomic DNA and thus resulting in some clustering variation for some strains. CBS6093 belongs to the H2 lineage based on dendrogram B, but clusters outside any of the other lineages in dendrogram A, suggesting genomic deviation from other H2 strains, a finding also supported by mating type, β-glucosidase activity, and MALDI-TOF data. CBS7982 clusters together with other P1 strains as expected in dendrogram A, but clusters close to H2 strains in dendrogram B. Interestingly, CBS7982 was found to contain a mitochondrial sequence different from other P1 strains (CBS9595, CD866, and CBS9574, Supplementary Figure 1B).

**Table 1.** Strains used in this study with information about their lineage, source, country of isolation, NGS sequencing data analysis, usage for PFGE, AFLP, sanger sequencing, mating-type, FACS, LM, MALDI-TOF and physiology analyses.

| Lineage | Strain | Source | Country | Data availability |  | PFGE | AFLP | Sanger sequencing<br>5 loci | Mating type | FACS | LM | MALDI-TOF | Physiology |
| --- | --- | --- | --- | --- | --- | --- | --- | --- | --- | --- | --- | --- | --- |
|  |  |  |  | NGS data | NGS strategy |  |  |  |  |  |  |  |  |
| P1 | CBS 9595 | back (man) | Greece | PRJNA732434 | Reference assembly P1 | - | + | + | + | + | + | + | + |
|  | CBS 9575 | back (man) | Greece | - | - | - | + | + | + | - | + | + | - |
|  | CBS 9574 | back (man) | Greece | PRJNA732434 | Read mapping | - | + | + | + | + | + | + | + |
|  | CBS 9589 | folds (man) | Greece | - | - | - | + | + | + | + | + | + | - |
|  | CBS 7982 | Skin of ear (man) | France | PRJNA286710* / PRJNA732434 | Read mapping | + | + | + | + | + | + | + | - |
|  | CD 866 | Poodle | Brazil | PRJNA732434 | Read mapping | - | + | + | + | + | + | + | + |
|  | CBS 9369 | Scalp of patient | Canada | - | - | + | - | + | + | - | + | + | + |
| P2 | CBS 8735 | bronchial wash | Canada | PRJNA732434 | Read mapping | + | + | + | + | + | + | + | + |
|  | CBS 14139 | urine, human | France | - | - | + | + | + | + | + | + | + | + |
|  | CBS 14141 | catheter, blood | France | PRJNA286710* / PRJNA732434 | Reference assembly P2 | + | + | + | + | + | + | + | + |
|  | PM 315 | anal swab neonate | Germany | PRJNA732434 | Read mapping | - | + | + | + | + | + | + | + |
| H1 | CBS 1878 NT | Dandruff, man | Unknown | PRJNA732434 | Read mapping | + | + | + | + | + | + | + | + |
|  | CBS 4171 | Ear of cow | Unknown | - | - | + | + | + | + | + | + | + | + |
|  | CBS 4172 | Skin of eland | Unknown | PRJNA286710* | Read mapping | + | + | + | + | + | + | + | + |
|  | CBS 7969 | Asian elephant | France | - | - | + | + | + | + | + | + | + | + |
|  | CBS 9365 | Elephant in zoo | France | PRJNA732434 | Read mapping | + | + | + | + | + | + | + | + |
|  | CBS 5101 | skin scales,<br>from PV | USA | - | - | - | + | + | + | - | + | + | - |
|  | CBS 4162 | Ear of pig | unknown | - | - | - | + | + | + | - | + | + | - |
| H2 | CBS 6000 | Dandruff | India | PRJNA732434 | Read mapping | + | + | + | + | + | + | + | + |
|  | CBS 6001 | PV | India | PRJNA732434 | Read mapping | + | + | + | + | - | + | + | - |
|  | CBS 6093 | Unknown | unknown | - | - | - | + | + | + | - | + | + | - |
|  | CBS 7019 NT | PV on trunk of<br>15-year-old girl | Finland | PRJNA732434 | Read mapping | + | + | + | + | + | + | + | + |

H1 = hybrid lineage 1, H2 = hybrid lineage 2, P1 = Parental lineage 1, P2 = parental lineage 2, PFGE = Pulsed-Field Gel Electrophoresis, AFLP
= Amplified Fragment Length Polymorphism. NT = neotype strain, + = analysis performed, - = analysis not performed, \*Data from Wu et al.

### Genome analyses confirm two independent hybridization events and their parental lineages

To confirm the existence of two independent hybridization events, 13 strains (Table 1) comprising representative isolates of all four AFLP-determined lineages were further compared in detail at the genomic level. To this end, we performed the genome assembly of the representative isolate of each of the parental lineages to serve as references for comparison with hybrid genome sequences (CBS9595 representing P1: 8.2 Mb, 8 scaffolds; and CBS14141 representing P2: 8.3 Mb, 9 scaffolds; Supplementary file 1A, Table 2, Materials and Methods for more details). The sequence alignment of these genome assemblies has shown that P1 and P2 lineages have an overall sequence similarity of 89 %. Therefore, a read mapping approach where all the reads are aligned on a single haplotype would not be possible, and hybrid sequencing reads were mapped to a combined reference including the P1 and the P2 (CBS14141: 8.3 Mb, 9 scaffolds) genome assemblies (Materials and Methods for more details) to identify the source and sequence of the different sub-genomes of the hybrids (Figure 2a).

**Figure 2.**
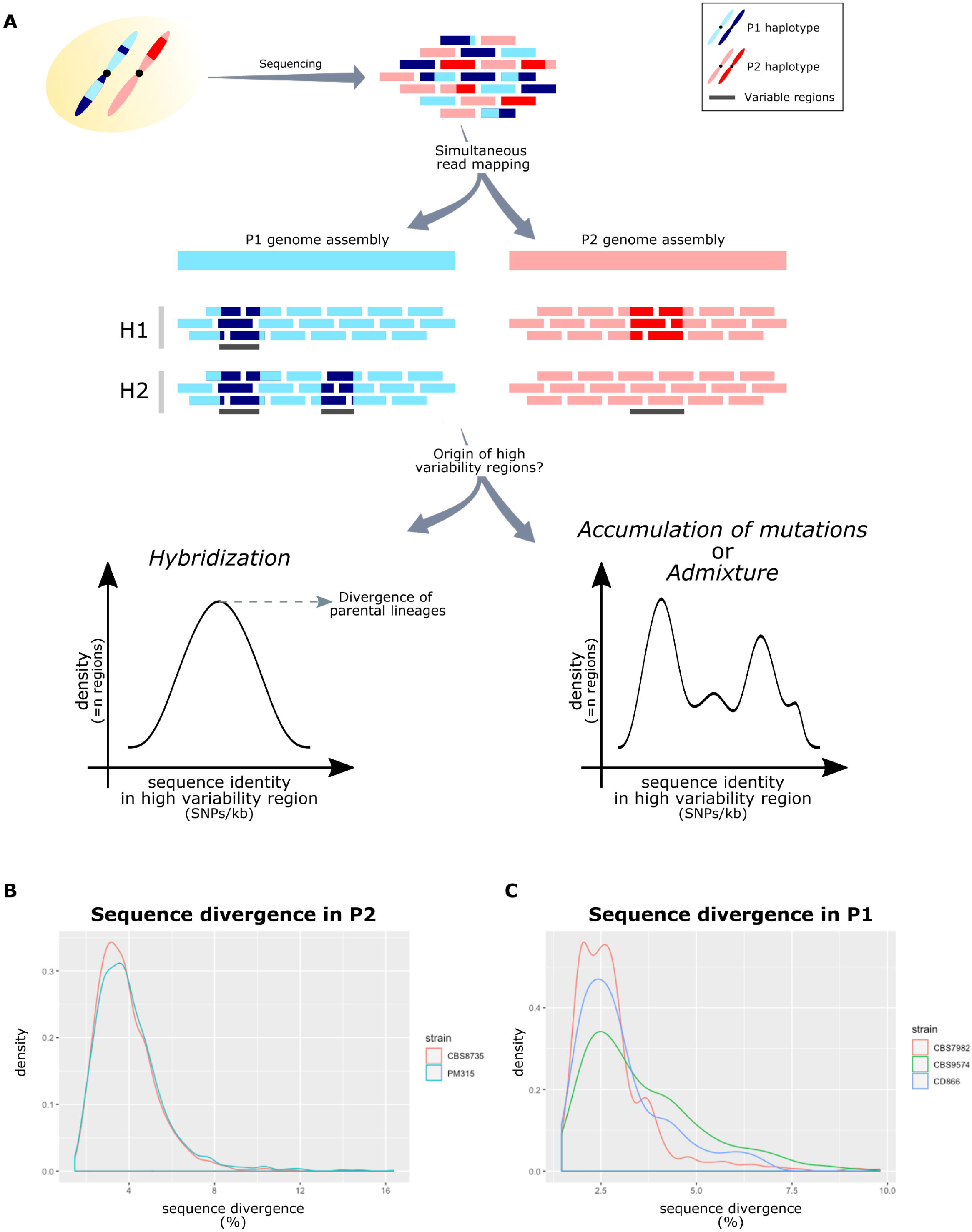
Analysis of the genomic patterns of hybrid genomes. **A)** Hybrid genomes were sequenced originating sequencing reads from P1 (blue rectangles) and P2 (pink rectangles). These reads were simultaneously aligned to the combined reference of P1 and P2. Light blue and light pink correspond to the alleles present in this reference. Dark blue and dark pink correspond to alleles which are aligned in P1 or P2, respectively, but present lower sequence identity forming blocks of genomic variability. Differences in the patterns of genomic variability were used to determine the different hybrid lineages. Estimated sequence divergence between the two alleles (i.e. between dark blue and light blue, or between dark pink and light pink) in terms of SNPs/bp in the blocks of genomic variability were used to determine the origin of such blocks: hybridization or admixture between different strains. **B)** Sequence divergence in the blocks of genomic variability of P2 lineages show a single density peak, suggesting a hybrid origin. **C)** Sequence divergence in the blocks of genomic variability of P1 lineages show multiple density peaks, with a single peak shared by all strains, not allowing the exclusion of any of the above-mentioned scenarios.

**Table 2.**
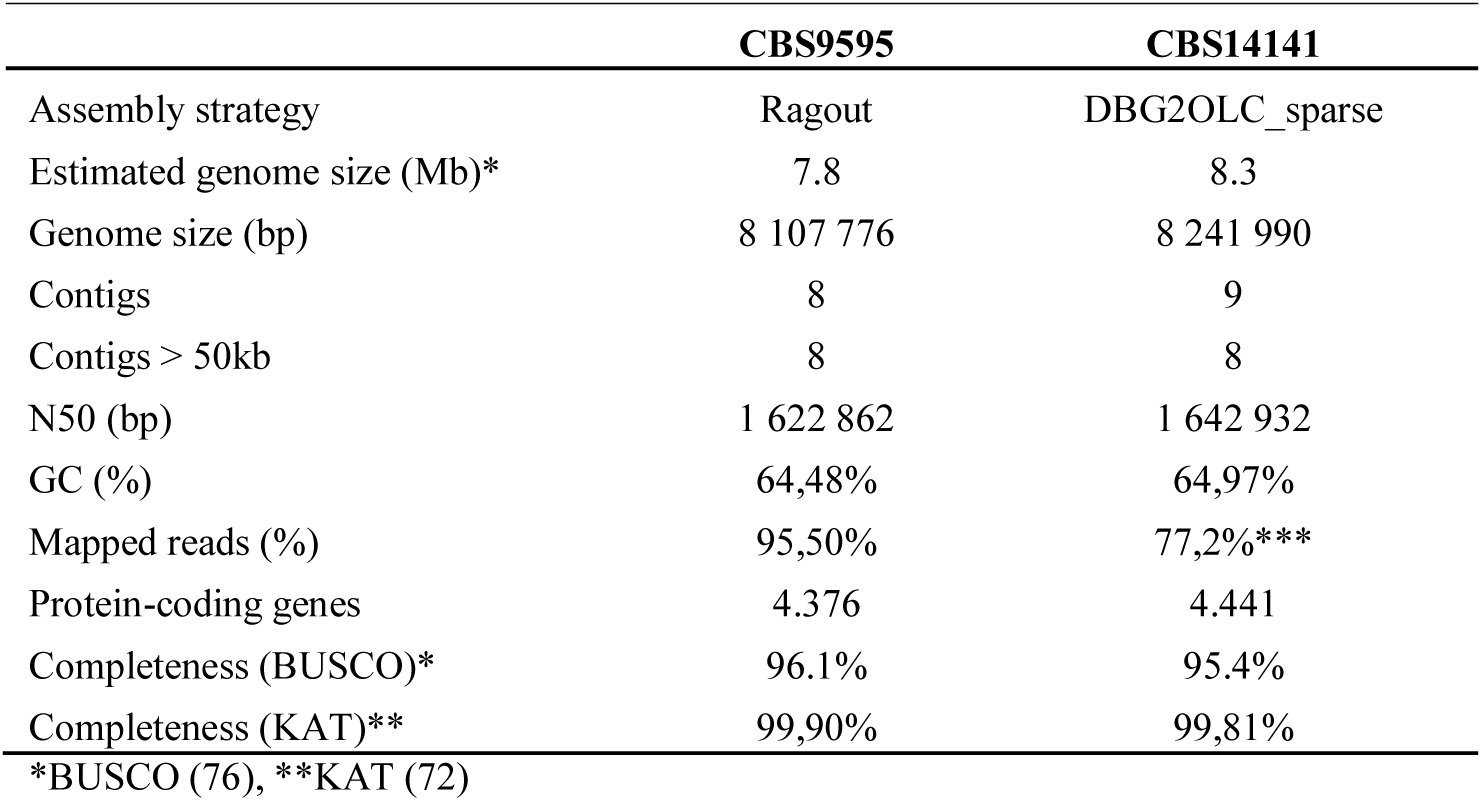
Summary of assembly statistics for the best genome assembly obtained for CBS9595 and CBS14141.

AFLP-determined H1 and H2 lineages on average presented 6 SNPs/kb (all homozygous given the alignment to a combined reference, Supplementary Table 1), with the majority of them (>85%) corresponding to P2 for all hybrid strains, thus suggesting that CBS9595 is possibly closer to the actual parent 1 than CBS14141 is to the actual parent 2. Noteworthy, these SNPs were not homogeneously distributed along the genome, but rather formed blocks of variants (see Supplementary Figure 1a and Materials and Methods for more details on block definition). Therefore, based on the assumption that strains originating from the same hybridization event would share the same blocks of variants, we utilized these as signatures of the hybrids’ evolutionary past and compared them between the different hybrids to confirm the number of hybridization events. Jaccard metrics revealed a high block overlap between the strains identified by AFLP as H1 (>84.6%), and the strains identified by AFLP as H2 (>91.8%), thus confirming the results of the AFLP analysis. It is important to highlight that among the H1 strains, CBS4172 presented the lowest similarity with its peers. While this strain revealed 84.6% and 84.9% block overlap with CBS1878 and CBS9365, respectively, the latter two strains had an overlap of 99%. The block overlap in strains of hybrid lineage H2 was more homogeneous. The overlap of the high polymorphic regions between strains of H1 and H2 varied between 20.8% and 23.1%, thus suggesting independent origins. Together, these results confirm that the hybrid lineages resulted from two independent hybridization events between P1 and P2. These findings suggest an apparent propensity of P1 and P2 to hybridize, and for their hybrids to survive.

### Genomic divergence uncovers the hybrid origin of P1 and P2

Considering the possible ancestry of the highly polymorphic blocks in the two hybrid lineages, we sought to determine their presence in the parental lineages. To this end, the same methodology was followed as for the hybrids (see Materials and Methods), and we verified that indeed, the strains of both P1 and P2 that were not used for genome assembly also harbor highly polymorphic genomic regions. A comparison between these blocks and the ones observed in the hybrid strains showed that the highest overlap between H1 strains and P1 and P2 is 14.6% and 21%, respectively. Similarly, comparing the regions of high variability in the hybrid lineage H2 with what was detected in P1 and P2 revealed that the highest overlaps were 64% and 13%, respectively. This is in line with our previous observations in the hybrid genomes and suggests that none of the strains sequenced for the P1 or P2 lineages represents the direct parental strain of the hybrids, but rather their close relatives.

The origin of these blocks of high genomic variability was unknown but, as reported by Mixão and Gabaldón (37) for *C. albicans*, two possible models could explain their existence: i) continuous admixture between different strains, and ii) a hybrid ancestor that experienced genomic recombination. A key aspect to distinguish these models is the divergence between the reference genome and the observed sequence, which we expect to vary between the different blocks in the first model, and to be similar among them in the second one (37). Therefore, we next assessed the level of divergence between these blocks and the reference genome in both lineages. For P2 strains, we identified a clear single peak of sequence divergence in both CBS8735 and PM315 (Figure 2b), suggesting that the genomic material of all the blocks was acquired at a single time-point, which in turns supports the existence of a hybrid ancestor. Our estimations point to a current haplotype divergence between the strains of parental lineage P2 of 4% to 4.5% (Supplementary Table 1). To determine if any sequenced strain could be the alternative parent of P2 lineage, we performed a BLASTn search in the NCBI genome database. The only hit obtained was the CBS14141 (the same strain we use as the P2 reference) genome with 95.2% sequence similarity, a value consistent with our estimations. This means that the alternative parent of the P2 lineage has not been sequenced thus far.

For the P1 lineage, this analysis was complicated by the limited number of polymorphic blocks (min 78 - max 463 blocks), and their short size (average = 245 bp), because a variation in a single polymorphism can have a large impact on the estimation of the sequence divergence. Indeed, despite the observed overlap between the peaks of sequence divergence in the different P1 strains, we also detected multiple sequence divergence peaks (Figure 2c). Therefore, we could not exclude any of the above-mentioned models (i.e., continuous admixture between different strains, or a hybrid ancestor that experienced genomic recombination) and properly estimate the sequence divergence. Even so, we performed a BLASTn search in the NCBI genome database using the longest blocks (>500 bp), and we verified that their best hits always presented a sequence similarity of ∼93% with the CBS14141 genome (P2 lineage), thus suggesting that P1 also has a hybrid origin.

### Mitochondrial genome analyses provide further proof for hybrid origin of parental lineages; all hybrids inherited mitochondria from P2

We next focused on comparing the mitochondrial genome sequences of the different *M. furfur* lineages. Genomic reads of each parental strain were mapped against mitochondrial genomes of the respective strains. As reference mitochondria for the P2 lineages, we selected the CBS14141 mitochondrial sequence available in NCBI (accession number: KY911086.1). As expected, all P2 strains shared the same mitochondrial genome. In the P1 lineage, the publicly available mitochondrial genome of CBS7982 (NCBI accession number: KY911085.1) was used as a reference for mapping sequencing reads from P1 strains. Mapping results revealed the presence of two different mitochondrial sequences in the P1 lineage with the mitochondrial sequence of CBS7982 differing from that of CBS9595, CD866, and CBS9574 (Supplementary Figure 1b). To compare both P1 mitochondrial types, a draft mitochondrial genome assembly with 49kb was generated for CBS9595 (see Materials and Methods). A BLASTn search with this mitochondrial genome against the NCBI nr database revealed that its sequence is equally distant to that of the mitochondrial genomes of CBS7982 (P1 lineage) and CBS14141 (P2 lineage), with a similarity of 93.45%. These results are in line with our observations for the polymorphic regions of the nuclear genome, further providing support that P1 results from the cross of two diverged lineages, i.e. hybridization, and CBS7982 harbors the mitochondrial genome of the alternative parent. A recent study assessing three mitochondrial genomic loci of 43 *M*. *furfur* strains, found two rather divergent mitochondrial clades for samples belonging to the P1 lineage (45), supporting our results and suggesting a wider presence of two different mitochondria among P1 strains. When reads of the strains belonging to both hybrid lineages were mapped against the mitochondrial reference genomes, it was observed that all hybrid strains inherited the mitochondria from the P2 lineage.

Together, these analyses suggest that both parental lineages are not genetically “pure” but are genomic mosaics likely resulting from a hybridization event between two unknown lineages that are approximately 7% divergent in their mitochondrial genome in the case of P1 (divergence in the nuclear genome could not be confidently estimated), and 4% divergent in their nuclear genome in the case of P2 (Figure 3). The observed small differences in the genetic mosaics detected in the nuclear genomes in each of the parental lineages suggest that strains in each lineage (P1 and P2) diverged before some of the recombination events occurred, a scenario which implies a non-haploid state of the strains at the time of their respective divergence. Nevertheless, all analyses performed in this study support a haploid state for all of them (Supplementary Figures 2 and 3A), indicating independent ploidy reduction.

**Figure 3.**
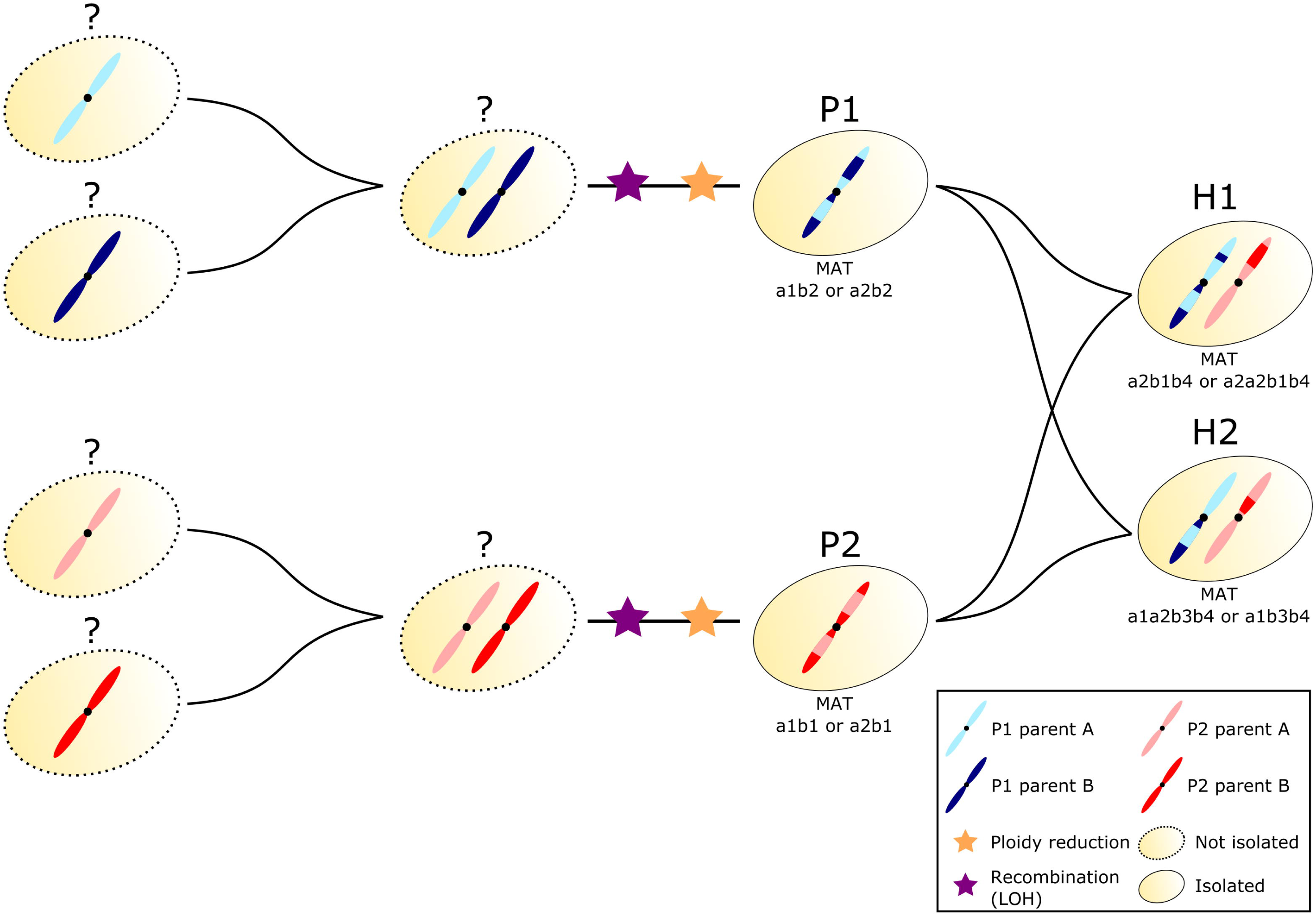
Comparative genomics analysis suggests multiple hybridization events in the evolutionary path of *M. furfur* lineages.

### *M*. *furfur* hybrid lineages are mostly diploid with some LOH

Karyotype data from this study and previous publications (46, 47) demonstrate significant chromosomal variation in *M. furfur* with hybrid strains containing additional chromosomal bands when compared to strains from their hypothesized haploid parental lineages (Supplementary Figure 2a). Moreover, FACS ploidy analysis for H1 strains displayed a DNA content between 1n and 2n suggesting aneuploidy, whereas the DNA-content of both analyzed H2 strains is consistent with assignment as diploids (Supplementary Figure 2b). Based on whole-genome sequencing data analysis, the analyzed strains for both hybrid lineages generally seem of diploid nature (Supplementary Figures 4), with a few exceptions, such as a putative triplication of a chromosome from P2 in the hybrid strain CBS9365 (H1, Supplementary Figure 4).

Considering the non-haploid state of these hybrid strains, we decided to look for LOH events, an important feature to restore genomic stability through the erasure of one of the haplotypes of the recombining region (34). In the case of the *M. furfur* hybrids analyzed in this study, we estimated that LOH covers approximately 20% of the genome (Supplementary Table 1). This is a low value when compared to other fungal hybrids, such as *Candida* hybrids, where LOH has been estimated to cover at least 50% of the genome (37–40, 44). Considering that sequence divergence between the parental lineages is higher in *M. furfur* (∼10%) than in *Candida* hybrids (∼4%) (37–40, 44), we hypothesize that the lower occurrence of LOH in *M. furfur* may be related to a lower number of potential recombining sites. Indeed, a recent study on *C. neoformans* x *C. gattii* hybrids, which present 7% sequence divergence, has shown that they experience fewer recombination events (41).

The direction of the LOH event, i.e. which allele is retained, varies from hybrid to hybrid, or even from niche to niche according to the most advantageous phenotypes (48). While in some hybrids there seems to be a tendency to retain the allele of a given parental lineage, in others this appears to be a random process (34, 40, 44, 49, 50). In *M. furfur* H1 and H2 hybrid lineages we found that >60% of the genome covered by LOH corresponds to the allele of P2, except for the strains CBS6001 and CBS7019 where this value is 45% (Supplementary Table 1). Although this result may suggest a slight tendency to retain the allele of P2 parental, we consider it to be not sufficiently clear or strong, as in other hybrids where >80% of the genome retains the allele of the same parent (49, 50). Indeed, if instead of focusing on the percentage of the genome covered by LOH, we analyze the number of LOH events that favored each of the parental alleles, we see that in three out of the six hybrid strains ∼56% of the events tended to P2, while in the other three this number is reduced to 43%, thus suggesting high stochasticity in the process.

### *M. furfur* possesses the genetic machinery of a pseudo-bipolar mating system

To understand the origin of the hybridization events leading to the H1 and H2 lineages, analysis of the mating-type genes of *M. furfur* was carried out (see Supplementary file 1B for a detailed description of the results). The *MAT* genes of *M. furfur* haploid strains were identified based on similarity searches with the *MAT* genes of two opposite mating-types of *M. sympodialis* (strain ATCC42132, *MAT a1b1*; strain ATCC44340, *MAT a2b2*). Two *MAT a* and two *MAT b* loci were identified in the haploid strains of *M. furfur*. The *MAT a* and *b* loci of the *M. furfur* haploid strains analyzed are ∼ 590 kb apart (Supplementary Table 2A), suggesting a pseudo-bipolar configuration. A representative comparison of the *MAT* loci of CBS14139 (*MAT a1b1*) and CBS7982 (*MAT a2b2*), as model strains, is shown in Figure 4. The *M*. *furfur MAT* structure reflects that of the *MAT* locus of *M. yamatoensis* (*16, 20*) and differs from the *M. globosa* and *M. sympodialis MAT* loci, for which *MAT a* and *MAT b* are ∼167 kb and ∼140 kb apart, respectively (20, 22). This is consistent with a whole-genome based clustering that groups *M. furfur* and *M. yamatoensis* in a separate phylogenetic cluster within the *Malassezia* genus (16). In all *M. furfur* haploid strains analyzed, the two *MAT a* locus genes are divergently oriented, corroborating previous findings for other *Malassezia* species (16, 20, 22). Moreover, a comparison between the *MAT a1* and *a2* loci revealed that the *Mfa* and *Pra* genes have an opposite arrangement albeit being located in highly syntenic flanking regions (Figure 4). In *Basidiomycetes*, the tetrapolar mating system is thought to be ancestral, and transition from a tetrapolar to bipolar system may be linked to the evolution of pathogenicity (51, 52). For example, in the genus *Cryptococcus*, pathogenic species have a bipolar mating system, whereas closely related non-pathogenic species such as *Cryptococcus amylolentus* have a tetrapolar mating system (53). Based on findings for the red yeast *Sporobolomyces salmonicolor* (cited as *Sporidiobolus salmonicolor*), the pseudo-bipolar mating system was proposed to be a gradual stage in the transition from a tetrapolar to bipolar system (52).

**Figure 4.**
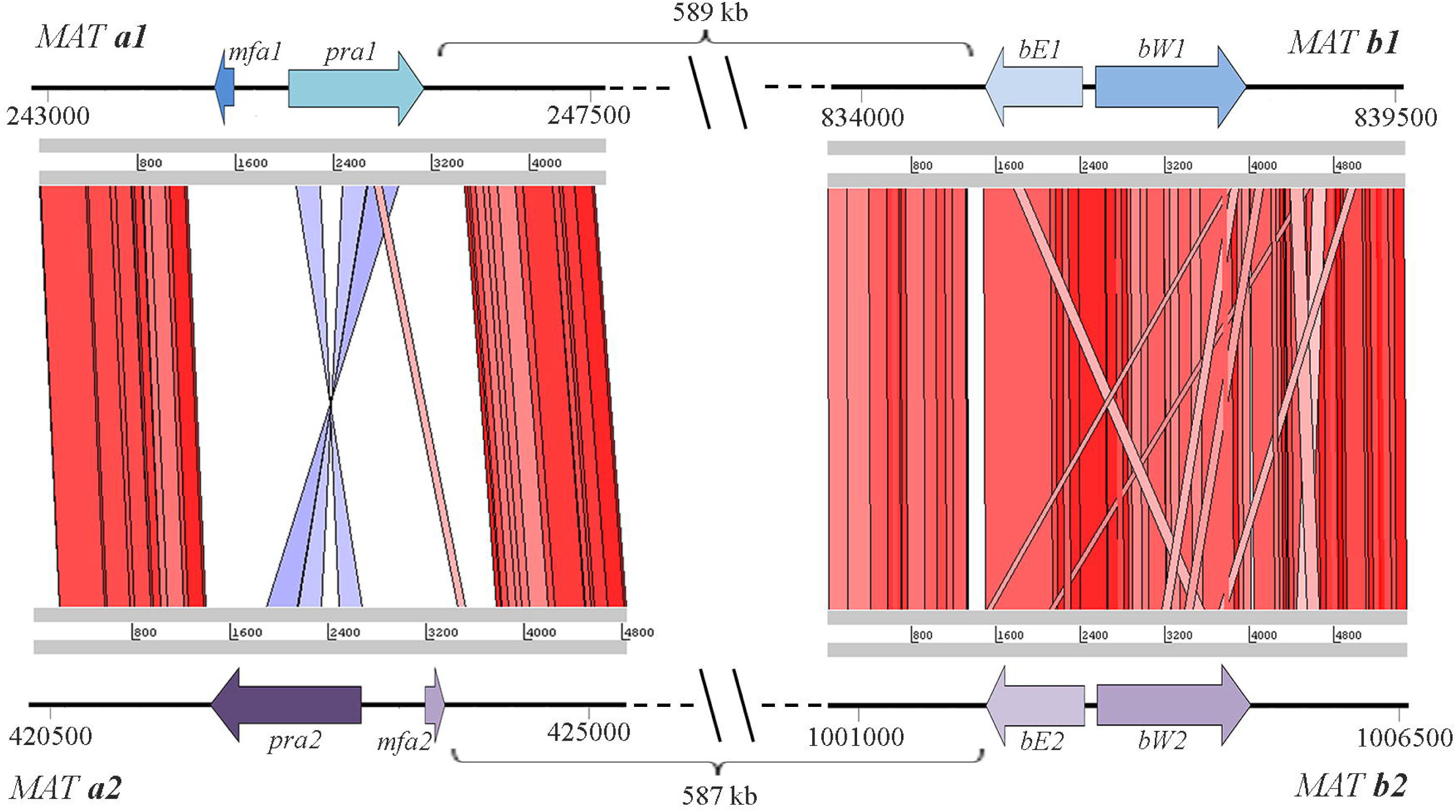
Schematic representation and comparison of mating loci for two mating compatible parental strains: *MAT* a1b1 loci of *M. furfur* strain CBS14139 (P2), and of the *MAT* a2b2 loci of *M. furfur* strain CBS7982 (P1). The coordinates of the genes in the genome scaffold are indicated. The two *MAT* regions were aligned with tBLASTx and visualized using ACT Artemis. The red and blue bars indicate regions of similarity, with red bars corresponding to regions of similar orientation and blue bars indicating regions oriented in opposite directions.

Hybrid genome searches with the *MAT* genes of haploid *M. furfur* identified two *MAT a* loci and two additional *MAT b* loci in the *M. furfur* hybrid strains CBS1878 (H1) and CBS7019 (H2). The sexual identity of the *b* loci among strains from all lineages was assigned following sequence comparison and phylogenetic analysis of the predicted proteins, resulting in *MAT b1*, *b2*, *b3*, and *b4* alleles, with *MAT* b3 and *MAT* b4 being present only in the hybrids, and closely related to *MAT* b1 and *MAT* b2, respectively (Figure 5C). In particular, based on genome data (both genome assembly and read mapping data), strain CBS1878 was designated as *MAT a2a2b1b4,* and strain CBS7019 as *MAT a1a2b3b4*. Interestingly, CBS1878 did not contain *MAT a1* but the *a2* locus was present in two copies. A closer inspection of the read alignment on the *MAT* locus with IGV (54)revealed that part of the Illumina paired-end reads flanking this duplication in a2 had their mate aligning in the edges of the region corresponding to the a1 allele (region present in the reference P1, but without read coverage in H1 strains), suggesting the occurrence of a LOH from an ancestral *a1a2* state to a derived *a2a2* configuration. Based on read mapping results, a similar occurrence seems to have happened for H1-strains CBS 4172 and CBS9365 (Supplementary Figure 1c).

**Figure 5.**
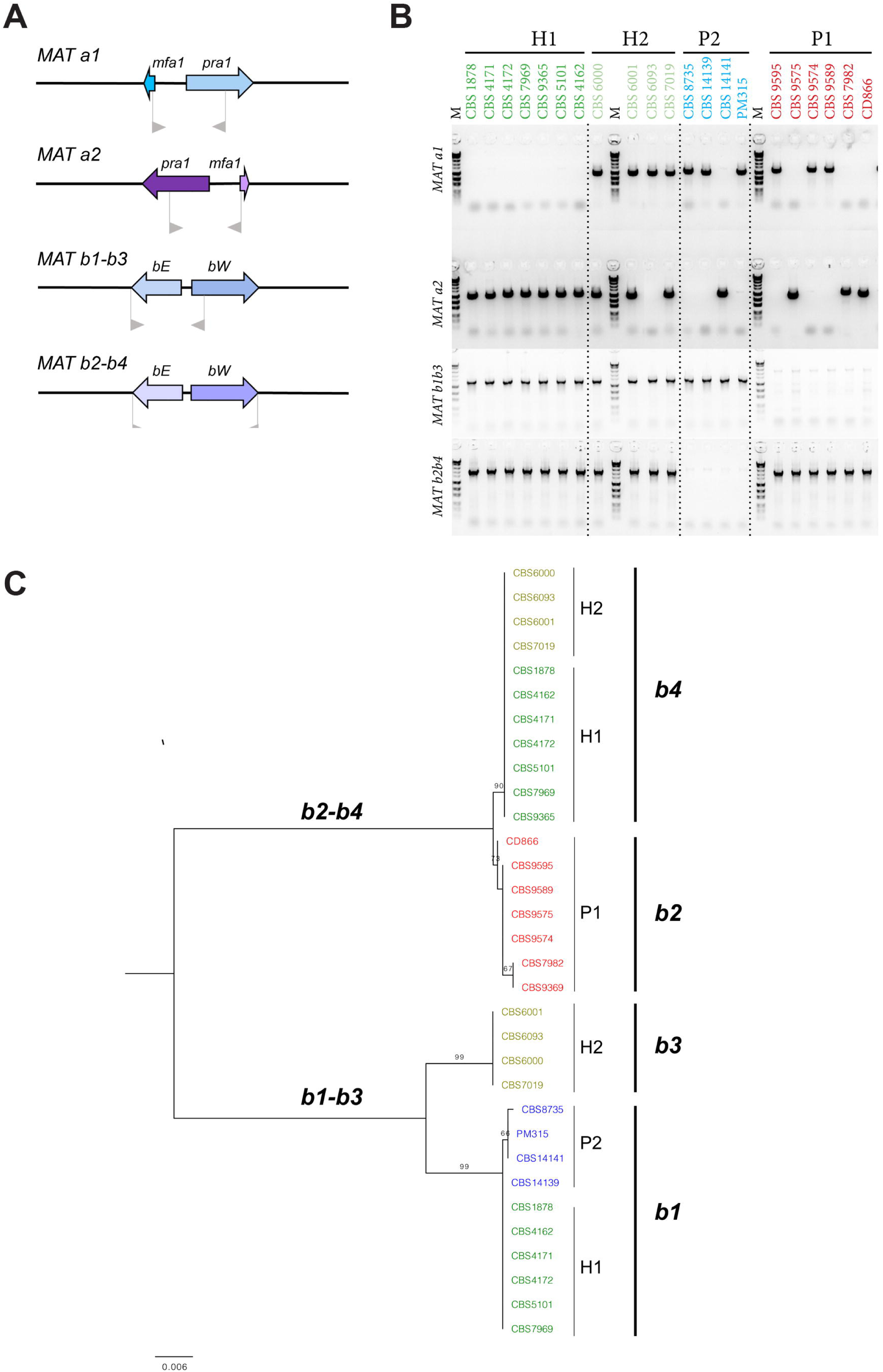
Mating typing assay results. A) Schematic presentation of primer positions in the MAT genes. B) Agarose gel electrophoresis picture, showing PCR results for all assessed strains for Mat a1, MAT a2, combined MAT b1-b3, and MAT b2-b4. To distinguish between b1 and b3, and between b2 and b4, the MAT B PCR-positive products require sequencing. C) Phylogenetic tree based on the Maximum Likelihood method and Tamura-Nei model with 500 bootstrap replications, representing the MAT B loci, resulting in four main clusters.

### A molecular assay for *M*. *furfur* mating type identifies mating-compatible strains in the parental lineages

Following these initial findings for the *M*. *furfur* mating-type regions, a PCR assay was developed with primers that specifically amplify *MAT a1*, *MAT a2*, *MAT b1-b3*, and *MAT b2-b4*. PCR results for the *MAT a1* and *a2*, and sequencing analysis of *MAT b1-b3* and *b2-b4*, indicated additional mating-type configurations (Figure 5A-C and Supplementary Table 2B). Both P1 and P2 parental lineages contain strains with either *MAT a1* or *MAT a2*, while all P1 strains are of the *MAT b2* type and all P2 strains are of the *MAT b1* type (Figure 5A-C and Supplementary Table 2B). The presence of *a1b1, a2b1, a1b2* and *a2b2 MAT* combinations supports the hypothesis that recombination occurs within this region, corroborating findings on the pseudo-bipolar *MAT* structure of *M. sympodialis* and in contrast with the lack of recombination reported for *Ustilago hordei*, which is bipolar (20). Our findings also highlight incompatibility at the B loci within each parental lineage, despite compatibility being present at the *MAT a* locus, which might explain why sexual reproduction could not be observed under laboratory conditions. Attempts to cross compatible *MAT a* and *MAT b* strains belonging to the P1 and P2 parental lineages were also carried out, but also in this case sexual reproduction could not be observed. It seems likely that two individual mating events between representatives of both parental lineages originally resulted in hybrid strains that contained the combination *MAT a1a2b1b2*, yet our data showed that various alterations of the mating loci occurred in the hybrids. All H1-strains lost the *MAT a1* copy and may have duplicated the *MAT a2* allele (only confirmed for CBS1878, CBS4172, and CBS9365). Strains belonging to the H2 hybrid lineage still have both parental *MAT a* copies, with the exception of CBS6093, that seems to have lost the parental *MAT a2* copy.

Additionally, two unique *MAT b* arrangements were observed in the hybrids: *MAT b4* in both hybrid lineages, with similarity to *MAT b2* of the P1 strains; and *MAT b3* which is only present in H2 strains, and is a phylogenetic sister of *MAT b1* of the P2 strains (Figure 5A-C and Supplementary Table 2B). Strains of hybrid lineage H1 retained the *MAT b1* copy. Considering that recombination in the *MAT* locus has previously been observed in other hybrid lineages (40, 55, 56), and is associated with a possible restoration of hybrid fertility (55, 56), we hypothesize that strains in lineages H1 and H2 may have undergone genomic changes leading to re-establishment of a viable sexual state.

### Targeted sequencing of five nuclear loci and genomic data suggest P1 and P2 lineages may be two separate species

The species *M. furfur* is represented by two neotype cultures, namely CBS1878 and CBS7019, corresponding to the respective names *Malassezia furfur* and *Pityrosporum ovale* (synonym of *M*. *furfur*), but these belong to the hybrid lineages H1 and H2, respectively. Many species are described based on a limited number of nuclear DNA loci, ribosomal loci ITS and LSU being two of the most frequently used taxonomic markers in fungi, but these loci may not reflect the genetic heterogeneity of a yeast strain sufficiently as is also shown in this study for the ITS of ribosomal DNA and three protein coding genes (β-tubulin, chitin synthase (*CHS2*), and translation elongation factor 1-α (EF1-α) (Supplementary Figure 3A1-5). Phylogenetic analysis of published sequencing data for the intergenic transcribed spacer (IGS) of the rDNA resulted in separate clusters for each of the studied lineages, making it a potential diagnostic tool for identifying hybrids belonging to H1 or H2 among genetically uncharacterized *M*. *furfur* isolates (Supplementary Figure 3A2). As chromatograms of hybrid strains for the protein coding genes possessed multiple sites with two different nucleotide peaks representing both parental backgrounds, they were phased into two sequences representing their respective parental copies (Supplementary Figure 3A3-5). Based on ITS data, both hybrid lineages H1 and H2 cannot be distinguished from the parental lineage P2, yet for the most part they contain genetic material from both P1 and P2. Type strains or neotype strains are reference strains, often used as representatives for a species in specific functional assays, but as in this case they represent hybrid lineages with a combined genomic content of two haploid parental lineages, they may not be the best choice to serve as reference strains and should probably be disqualified to serve as neotypes. At present, all strains considered in this study are classified as *M. furfur*. However, based on sequence divergence between the P1 and P2 lineages for the five analyzed nuclear genomic loci it is likely that P1 and P2 may represent two closely related species. For example, ITS sequence similarity between P1 strain CBS9595 and the P2 strain CBS14141 is 98% but their similarity for protein coding genes is much lower: 97.3% for EF1-α, 96.4% for β-tubulin, and 87.6 % for *CHS2* respectively. According to a study that assessed more than 9 000 yeast isolates to establish species and genus thresholds for ribosomal ITS and LSU regions, a species-threshold of 98.41% was presented for ITS (57), which supports the assignment of these lineages into two species. Furthermore, as above-mentioned, based on whole-genome sequence alignments for CBS9595 (P1) and CBS14141 (P2), an overall sequence similarity of 89 % was estimated, further supporting that these strains represent two species.

### Hybrid lineages H1 and H2 show different morphologies but limited differentiation based on traditional physiological properties

At the phenotypic level, cells belonging to hybrid lineages differ in size and shape from parental lineage cells (Figure 6, Supplementary Table 3), with hybrid cells being thinner and more elongated. However, growth profiling experiments traditionally used for *Malassezia* species identification, seem not to differ significantly between hybrid and parental lineages (Supplementary Table 4). One noteworthy observation is that all P2 strains were positive for β-glucosidase activity and strains from the other lineages were all negative, except for one P1 strain (CBS9589) and for one H2-strain, CBS6093. This latter strain also seems somewhat different from other H2-strains based on mating type, AFLP-pattern, and Matrix-Assisted Laser Desorption Ionization-time Of Flight (MALDI-TOF) mass spectrum (Supplementary Figure 3B) and would be an interesting candidate for further genomic exploration. Variable results for Cremophor EL utilization and β-glucosidase activity have been described previously for *M*. *furfur* isolates (12, 15, 23–26) and may be an expression of the heterogeneity of the species. Analysis of mass spectra generated with a Bruker MALDI Biotyper also illustrated the differences between P1 and P2 lineages and formed separate dendrogram clusters for H1, H2, P1, and P2, with a few exceptions (Supplementary Figure 3B). The putative H2 strain CBS6093 clustered basal to clusters for P2 and H2, supporting above mentioned deviating findings for that strain. In addition, the MALDI-TOF mass spectrum of H1-strain CBS4172 clusters with the P2 strains. This divergent character is in line with a deviating PFGE chromosomal banding pattern and this strain has lost *MAT* a1, and only has one copy of *MAT* a2. These findings need to be repeated but confirm the current high heterogeneity present in the species *M*. *furfur* that might be better interpreted as a species complex.

**Figure 6.**
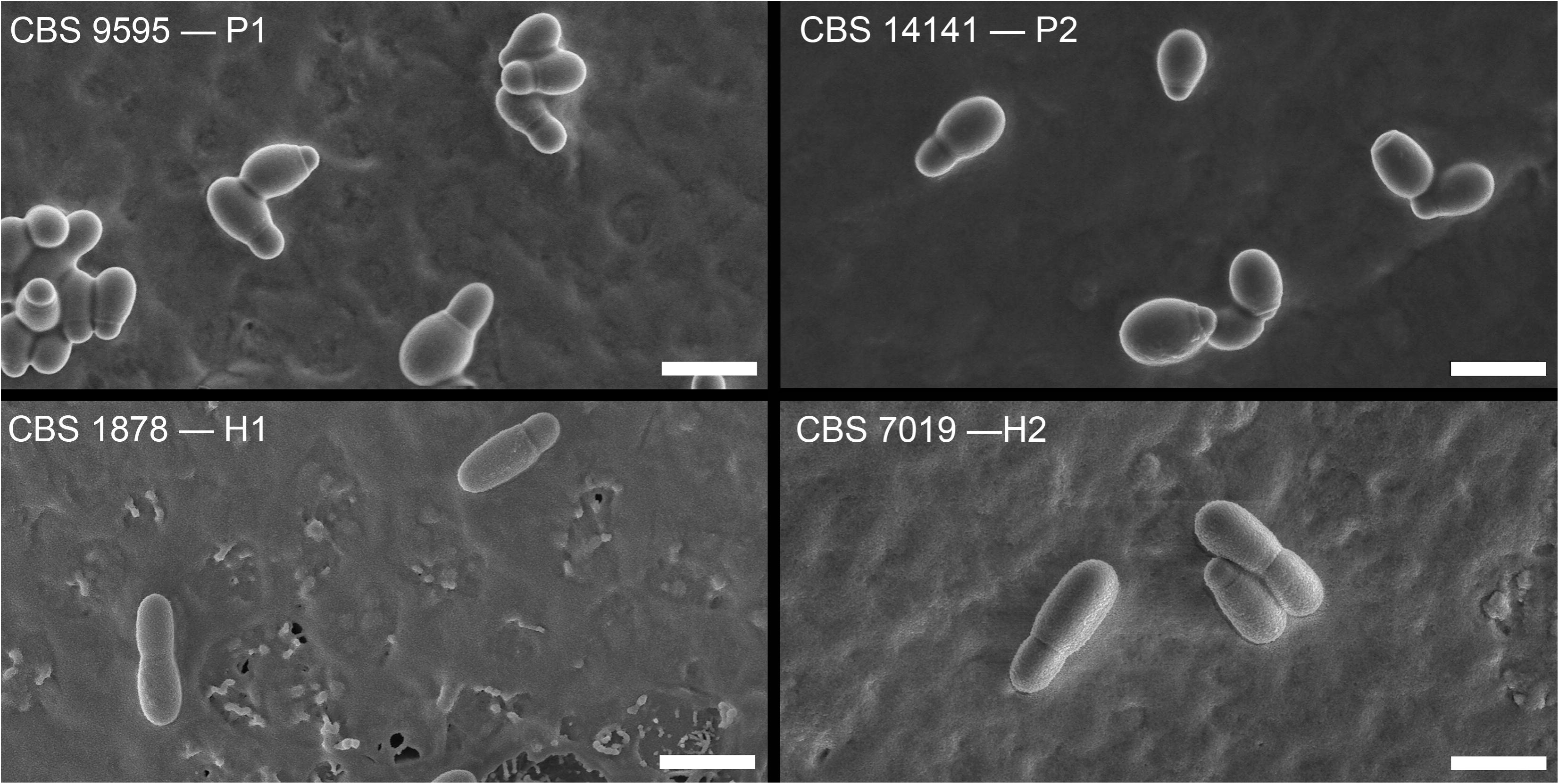
SEM photo plate for representative strains of each lineage, showcasing different morphologies between haploid parental strains and hybrid strains. Additional phenotypic data is available in Supplementary tables 3 and 4 for cell size measurements and traditional physiological data respectively. Bar = 5 μm

Based on AFLP, mating type analysis, and Sanger sequencing data, the H1 hybrid lineage consists of seven strains, five originating from animal skin and two from diseased human skin. Hybrid lineage H2 consists of four strains, all isolated from human diseased skin, with the exception of CBS6093 for which the origin is unknown. Of note, as has also previously been mentioned (30, 58), deep-seated isolates (e.g. blood, urine, rectal swabs) seem to almost exclusively belong to the genotype representing the P2 lineage. Such an observation could, at first sight, be at odds with the previously proposed hypothesis that hybridization plays a role in the emergence of pathogenicity (34). However, it is important to note that such a hypothesis does not imply that hybridization is the only mechanism leading to pathogenicity, and, in the particular case of M. furfur P2 lineage, our results show that it was likely to result from a hybridization event as well. Leong and colleagues explored antifungal susceptibility patterns of 26 *M*. *furfur* strains, including some strains also considered in this study (59). Four hybrid strains (H1: CBS1878, CBS7019; H2:CBS6000, CBS600) showed reduced susceptibility to certain azoles. This feature was not exclusive for strains from hybrid lineages H1 and H2 but rather seemed linked to disease state backgrounds of strains, and reduced azole susceptibility also included disease isolates belonging to parental lineage P2 (59). Whether hybridization may have facilitated genetic changes driving this reduced azole susceptibility in any of the lineages H1, H2, or P2, or whether this was the result from mere exposure to these drugs, remains to be elucidated. Considering that five out of the seven H1-strains are derived from animals, we hypothesize that the hybridization event for H1 may have facilitated a host-shift event between humans and animals. This hypothesis should be tested in future analyses including a larger sampling of strains.

In summary, this study identified two individual hybrid lineages H1 and H2, with P1 and P2 representing their parental lineages, although not the exact parental strains. We propose that the diploid hybrid lineages H1 and H2 are the result of two separate mating events between mating-compatible strains from the P1 and P2 groups. Interestingly, our genome analysis shows that both haploid parental lineages were themselves the result of prior hybridization events but became haploid before some of their members mated again to form hybrid lineages H1 and H2. The various hybridization events and subsequent further evolutionary changes have contributed to additional genetic diversification in *M. furfur*, but also at the phenotypic level, the hybrid strains differ from their parental lineages. The implications of possible hybridization-driven changes in pathogenicity, or adaptation to new environments, will drive further analysis and in depth examination comparing pathogenicity factors, lipid production and utilization, and salient physiological and phenotypic facets.

## Materials and Methods

### DNA extraction

To extract DNA for whole-genome sequencing, cells were grown on modified Dixon agar (mDA) (60) for 48 h at 30°C and harvested into 50 ml tubes. Yeast cells were lysed, using the QIAGEN Genomic DNA Purification procedure for yeast samples (Qiagen, Hilden, Germany), with minor modifications. Lyticase incubation was performed for 2 h at 30°C, and RNase/Proteinase incubation was performed for 2 h at 55°C. Genomic DNA was purified using Genomic-tip 100/G prep columns, according to the manufacturer’s handbook. For AFLP, MLST, and the mating type assay, gDNA extraction was performed following the CTAB method as described by (61) with the following modification: DNA was purified in two steps-first with phenol-chloroform and secondly with chloroform only.

### AFLP

AFLP analysis was performed according to (62) with some modifications. Various restriction enzymes and adaptor pairs were combined with multiple primer combinations. For each combination, a combined restriction ligation reaction was performed for 2 h at 37°C. AFLP combination A constituted of adaptor pairs MseI-a1(5’-GACGATGAGTCCTGAC-3’) with MseI-2a (5’-TAGTCAGGACTCAT-3’), and EcoRI_Adaptor1 (5’-CTC GTAGACTGCGTACC-3’) with HpyCH4IV (5’-CGGGTACGCAGTC-3’). A pre-selective PCR was performed with primers HpyCH4IV_core (5’-GTAGACTGCGTACCCGT-3’) and MseI_core (5’-GATGAGTCCTGACTAA-3’) for 20 cycles with annealing at 56°C, and a subsequent selective PCR was executed with primer pair HpyCH4IV_selectC_FAM (5’-/56-FAM/GTAGACTGCGTACCCGTC-3’) and MseI_selectTGAG* (5’-GATGAGTCCTGACTAATGAG-3’), for 10 cycles with an annealing temperature reducing from 66°C to 56°C, followed by 20 cycles with annealing at 56°C. AFLP combination B constituted of adaptor pairs MseI-1a (5’-GACGATGAGTCCTGA G-3’) with MseI-2a (5’-TACTCAGGACTCAT-3’) and EcoRI-1 with EcoRI-2 (5’-AATTGGTACGCAGTCTAC-3’, and was directly followed by a selective PCR reaction with primer pair EcoRI_selectA_FAM (5’-/56-FAM/GACTGCGTACCAATTCA-3’) and MseI_selectG (5’-GATGAGTCCTGAGTAAG-3’), for 20 cycles with an annealing temperature reducing from 66°C to 56°C, and additional 30 cycles with annealing at 56°C. PCR products were diluted 200x after purification and then combined with Orange600 size standard (Nimagen, Nijmegen, the Netherlands) before fragment analysis on a 3730xl DNA Analyzer (Thermo Fisher Scientific, Waltham, Massachusetts, USA). Data was imported and analyzed using Bionumerics v.7.6.3 (Applied Maths, Sint-Martens-Latem, Belgium), and dendrograms were created with UPGMA clustering and Pearson correlation coefficient. After purification of PCR products, all reactions were diluted 200x.

### Whole-genome sequencing

Thirteen samples were selected for whole-genome sequencing data analysis (Table 1). For three of them (CBS7982, CBS14141 and CBS4172), we retrieved Illumina data from SRA (Table 1) (16). For the remaining ones, whole-genome sequencing was performed at the Genomics Unit from the Centre for Genomic Regulation (group 1: CBS8735, PM315, CBS1878 and CBS7019) and at the Genome Institute Singapore (group 2: CBS9595, CD866, CBS9574, CBS6000 and CBS6001). Except when specified, the protocol was similar in both groups of strains. Libraries were prepared using the NEBNext Ultra DNA Library Prep kit for Illumina (New England BioLabs, United States) according to manufacturer’s instructions. All reagents subsequently mentioned are from the NEBNext Ultra DNA Library Prep kit for Illumina, if not specified otherwise. 1 μg of gDNA was fragmented by ultrasonic acoustic energy in Covaris to a size of ∼600 bp in group 1 and ∼300-400 bp in group 2. After shearing, the ends of the DNA fragments were blunted with the End Prep Enzyme Mix, and then NEBNext Adaptors for Illumina were ligated using the Blunt/TA Ligase Master Mix. The adaptor-ligated DNA was cleaned-up using the MinElute PCR Purification kit (Qiagen, Germany) and a further size selection step was performed using an agarose gel. Size-selected DNA was then purified using the QIAgen Gel Extraction Kit with MinElute columns (Qiagen) and library amplification was performed by PCR with the NEBNext Q5 Hot Start 2X PCR Master Mix and index primers (12–15 cycles in group 1, 6 cycles in group 2). A purification step was done using AMPure XP Beads (Agentcourt, United States). The final library was analyzed using Agilent DNA 1000 chip (Agilent) to estimate the quantity and check size distribution, and it was then quantified by qPCR using the KAPA Library Quantification Kit (KapaBiosystems, United States) prior to amplification with Illumina’s cBot.

Libraries were loaded and sequenced 2 x 125 on Illumina’s HiSeq2500 for group 1 and 2 x 101 on Illumina’s HiSeq2000 for group 2. Base-calling was performed using Illumina pipeline software. De-convolution was performed using the CASAVA software (Illumina, United States).

Four samples (CBS7982, CBS9595, CBS14141 and CBS1878) were additionally sequenced with PacBio long-read sequencing strategy. In addition, previously generated PacBio sequencing data was used for mating type analysis only, for five samples (CBS14139, CBS8735, PM315, CBS4172, CBS9369 and CBS7019). All samples were sequenced on the PacBio RSII platform (Pacific Biosciences, United States), and, except for CBS9595, libraries were prepared in 2015 with the DNA Template Prep Kit 3.0, and polymerase/template complexes were subsequently formed using Polymerase Binding Kit P6 v2, and then sequenced with Sequencing Reagent Kit 4.0. Sample CBS9595 was sequenced more recently, with the following specifications: the library was prepared using the SMRTbell Template Prep Kit 1.0, polymerase/template complexes were generated with DNA/Polymerase Binding Kit P6 v2, and the sample was sequenced using DNA Sequencing Reagent Kit 4.0 v2 with 360 min runtime per SMRTcell.

### *De novo* genome assembly

The genomes of the samples exclusively sequenced with PacBio (CBS14139, CBS8735, PM315, CBS4172, CBS9369 and CBS7019) were assembled with HGAP3 within the SMRT portal of PacBio, SMRTanalysis v3.1, with standard settings. The genomes of the four samples with short- and long-read sequencing libraries (CBS7982, CBS9595, CBS14141 and CBS1878) were assembled with a pipeline that combines short- and long-read assemblers. Briefly, Illumina reads were filtered and trimmed with Trimmomatic v0.36 (63) and assembled with Platanus v1.2.4 (64). PacBio reads were corrected with Canu (65) and assembled with DBG2OLC (v20180222) (66) using Platanus assembly, MaSurCA v3.3.0 (67), and WTDBG2 v2.1 (68). Ragout v2.2 (69) was used for scaffolding using DBG2OLC, WTDBG2 and MaSurCA assemblies. Assembly correction was performed with Pilon v1.22 (70). The assemblies’ quality was assessed with Quast v4.5 (71) and K-mer Analysis Toolkit v2.4.1 (KAT, (72)). The best assembly for each sample was chosen based on N50, level of fragmentation and estimated assembly completeness by KAT (72). Augustus Web-server (73, 74) was used for genome annotation, using *Malassezia restricta* proteome as training set (accession number: GCA_003290485.1 (75)). Predicted protein-coding genes completeness was assessed with BUSCO v4 using the Basidiomycota database (76). Functional annotation was performed with eggNOG-mapper web-server using the default settings (77).

### Read mapping and variant calling

All paired-end Illumina libraries were inspected with FastQC v0.11.5 (http://www.bioinformatics.babraham.ac.uk/projects/fastqc/) and trimmed and filtered with Trimmomatic v0.36 (63). Read mapping was performed with sppIDer pipeline (78) using a combined reference including the genome assembly of CBS9595 (as representative of P1 lineage, Supplementary File 1A for more details) and CBS14141 (as representative of P2 lineage, Supplementary File 1A for more details). To guarantee the proper correspondence between the scaffolds of both parentals, we aligned both genomes with the nucmer tool of MUMmer v3 (79). Average genome coverage for each sample was estimated with Samtools v1.9 (80). Variant calling was performed with HaploTypo v1.0.1 (81) selecting Freebayes v1.3.2 (82) as variant caller, and using the default settings for the remaining parameters. Read alignment was inspected with the Integrative Genomics Viewer (IGV) (54).

### Definition of blocks of high genomic variability

To determine for each of the haplotypes of the hybrid and the parental genomes the regions with high variability when compared to the reference, we used the methodology developed and tested by (40) for LOH blocks definition. Briefly, we used bedtools merge (83) with a distance of 100 bp to merge the homozygous SNPs of each sample, and we set a minimum polymorphic region size of 100 bp. These blocks were compared among the different strains using bedtools jaccard (83). The sequence divergence between the reference alleles and the allele of the polymorphic regions was calculated by dividing the number of SNPs overlapping such regions by the total number of base pairs covered by them. Of note, these regions did not represent LOH regions in the hybrids, as they only reflect the differences with the reference. Nevertheless, they were used to infer the patterns of the respective parental lineages.

### LOH blocks definition in the hybrid strains

As read mapping of the hybrid strains was performed simultaneously in both parental lineages, it was not possible to define LOH blocks based on the distribution of heterozygous variants, as usually performed (37, 40, 43, 44). Instead, an alternative approach where regions deleted in one parental and duplicated in the alternative one were used as an indicator of recombination, i.e. LOH. Therefore, LOH block definition in hybrid strains relied on read depth of coverage. Briefly, bedtools genomecov (83) was used to determine the number of reads covering each position. Positions covered by 0 reads were considered deleted, while those covered by 150% of the average genome coverage were considered duplicated. Similarly to the procedure developed by (40), we determined a minimum block size of 100 bp. Bedtools intersect was used to determine the intersection of the deleted regions of P1 and the duplicated regions of P2, and *vice-versa*. Only duplicated regions in one parent that intersect a deleted region in the alternative one were considered as LOH. An enrichment analysis of the genes overlapping LOH blocks was performed with FatiGO (84).

### Mitochondrial genome assembly

*De novo* genome assembly of CBS9595 mitochondrial genome was performed with NOVOPlasty v2.7.2 (85), using CBS7982 *cox2* sequence as seed (accession number: KY911085.1).

### Identification of *M*. *furfur* mating type region

P/R (*MAT A*) and HD (*MAT B*) loci of *M. sympodialis* strains ATCC42132 (*MAT a1b1*) and ATCC44340 (*MAT a2b2*) (20) were used as query for tBLASTn analysis on the PacBio genomic assemblies of *M. furfur* haploid strains. The designation of *M. furfur MAT A* loci was assigned following that of the closest *M. sympodialis* orthologs based on the *E-*value of the tBLASTn outcome, whereas that of *MAT B* loci was assigned according to phylogenetic clustering of the predicted concatenated HD proteins (see below). In all cases, the *MAT* genes identified in *M. furfur* strains were confirmed by reciprocal BLASTx on GenBank. The nomenclature of the *M. furfur MAT* genes follows that of the closely-related Ustilaginomycotina: *mfa* is the pheromone-encoding gene, *pra* is the pheromone receptor, *bE* and *bW* are the HD transcription factors, followed by a number to distinguish from different alleles (18).

Open reading frames of the *MAT* genes were predicted by comparison with their respective orthologs through BLASTx on GenBank, and using RNAseq data available for *M. furfur* CBS14141 (Bioproject PRJNA741845 (59)). DNA sequences of the *MAT* genes were aligned with MUSCLE (86) and their phylogenetic reconstruction was performed with MEGA7 (87) using the maximum likelihood method [Tamura 3-parameter model with Gamma distribution] and 100 bootstrap replications. Similarly, translated HD proteins were predicted with ExPASy translate tool (88), and concatenated bE-bW sequences were aligned with MUSCLE (86) and the respective maximum likelihood tree [Jones-Tailor-Thornton (JTT), uniform rates] with 100 bootstrap replications was obtained with MEGA7 (87).

The identified genes of the *MAT a1, a2, b1,* and *b2* loci of the *M. furfur* haploid strains were used as queries for BLASTn and tBLASTx analyses to identify the *MAT* regions in the *M. furfur* hybrid strains CBS1878, CBS4172, and CBS7019. The *MAT a1* and *MAT a2* designation followed that of the haploid strains used as input. For the identified *MAT B* loci, the DNA sequences of *bE* and *bW* genes and their predicted encoded proteins were aligned with MUSCLE (86) and then subjected to phylogenetic analysis with MEGA7 (87) as described above (data not shown).

### Characterization of *M*. *furfur* mating type

Primers for the amplification of *MAT a1, a2, b1,* and *b2* alleles were designed on the basis of an alignment of all *MAT* loci of the available *M. furfur* parental and hybrid strains derived from their genome sequences. The primers for *MAT A* loci specifically amplified the *a1* (JOHE44273, 5’-TTGGCAGAGTTGACAGGCT-3’; and JOHE44272, 5’-AACCATCCATGCTGACATTT-3’) or *a2* allele (JOHE44274, 5’-GAGCCACAAGATAATGTCAA-3’; and JOHE44275, 5’-AGACTTCCTGAACAGTGTCC-3’), while the primers for the *MAT B* loci amplified *b1* and *b3* (JOHE44491, 5’-TTCGGTTGACGGTCCCTCGGC-3’; and JOHE44492, 5’-ACCGCGACTGCGCATCCGCG-3’), or *b2* and *b4* (JOHE44494, 5’-TTCGCCAAATGTGTTCGGCC-3’; and JOHE44496, 5’-CAGCAACACCCGCCTCGCTT-3’). For the amplification of the *MAT A* loci, ExTaq (Takara) was used following the manufacturer’s instructions with the following PCR conditions: initial denaturation 2 min at 94°C, followed by 33 cycles of 30 sec at 94°C, 30 sec annealing at 58°C, 1 min 15 sec extension at 72°C, and a final extension of 5 min at 72°C. For the amplification of the *MAT b1-b3,* LATaq (Taqara) supplemented with GC Buffer II was used following the manufacturer’s instruction, and PCR conditions were: initial denaturation 2 min at 94°C, followed by 33 cycles of 30 sec at 94°C, 30 sec annealing at 60°C, 2 min extension at 72°C, and a final extension of 5 min at 72°C. The *MAT b2-b4* alleles were amplified using the same conditions as reported for *MAT b1-b3*, except for the use of LATaq (Taqara) supplemented with GC Buffer I. *MAT B* loci are high in G+C (∼65 %) and the use of specific Taq polymerase was important for their successful amplification.

The amplified *MAT* b1-b3 and *MAT* b2-b4 alleles were then sequenced using the primers used for amplification, and sequences were aligned using MEGAX with MUSCLE and then subjected to phylogenetic analysis using the maximum likelihood method [Tamura-Nei model] and 500 bootstrap replications (86, 89).

### Scanning Electron Microscopy

Hybrid- and parent strains were cultivated on modified Dixon (mDixon) medium for 72 h and a loop of cells was suspended in water. The cells were briefly vortexed to dislodge from each other. Droplets of 1, 2 and 3 µl were gently placed on mDixon agar and dried for 1 h in a laminar flow cabinet to fix the cells onto the agar. After pre-examination under a stereo microscope, small 4 x 4 mm selections with both individual cells and cells grouped together were cut out using a surgical blade (Swann-Morton, no. 11, Sheffield, UK) and glued on a copper sample cup with a small droplet of frozen-tissue medium (KP-Cryoblock, Klinipath, Duiven, The Netherlands) and subsequently snap-frozen in nitrogen slush, and transferred into an Oxford CT1500 Cryostation connected to a JEOL 5600LV scanning electron microscope (JEOL, Tokyo, Japan). Samples were sputter-coated (3 x 1 min) using a gold target in the cryostation. Electron micrographs were taken at an acceleration voltage of 5 kV.

### Data availability

Sequencing data, genome assemblies and annotations are available at NCBI database under the BioProject accessions PRJNA732434 and PRJNA779728.

## Acknowledgements

The authors thank Timothy James for reviewing our manuscript, Bart Kraak for some exploratory PCR and microscopy work, Simon Denil for helping with initial bioinformatics assessment of strain CBS1878, Claudia Cafarchia for providing strain CD866 and Marina Marcet-Houben for all the helpful discussions on the bioinformatics analyses. This work was supported by the European Union’s Horizon 2020 research and innovation programme under the Marie Sklodowska-Curie grant agreement No H2020-MSCA-ITN-2014-642095. TG group also acknowledges support from the Spanish Ministry of Economy, Industry, and Competitiveness (MEIC) for the EMBL partnership, and grants ‘Centro de Excelencia Severo Ochoa 2013-2017’ SEV-2012-0208, and BFU2015-67107 co-founded by European Regional Development Fund (ERDF); from the CERCA Programme / Generalitat de Catalunya; from the Catalan Research Agency (AGAUR) SGR857, and grants from the European Union’s Horizon 2020 research and innovation programme under the grant agreement ERC-2016-724173. TG also receives support from an INB Grant (PT17/0009/0023 - ISCIII-SGEFI/ERDF). G.I and J.H. were supported by NIH/NIAID R37 award AI39115-24 and R01 award AI50113-16A1. J.H. is fellow and co-director of the CIFAR program Fungal Kingdom: Threats and Opportunities. TD was supported by the A*STAR Industry Alignment Fund (H18/01/a0/016).

## Supplementary material

**Supplementary Table 1.** Summary metrics of the whole-genome sequencing analyses of M. furfur strains.

**Supplementary Table 2.** A) tBLASTn E-values for the identification of M. furfur mating type analysis and B) mating type PCR amplification results for studied strains.

**Supplementary Table 3.** Cell measurements with light microscopy after 72h at 33°C incubation on mDixon medium.

**Supplementary Table 4.** Temperature growth profiles and physiological characteristics of M. furfur strains. Growth at 6°C, 15°C, 24°C, 30°C, 33°C, 40°C, and 42°C on mDA (modified Dixon Agar), SGA (Sabouraud glucose agar), and YNBA (nitrogen base agar); Catalase activity; β-glucosidase activity; Tween diffusion test on SGA and YNBA at 15°C, 33°C, and 37°C for selected strains with Tween 20, 40, 60, 80 and CrEL (Cremophor EL).

**Supplementary File 1.** A) Genome assembly of *M. furfur* parental lineages and comparison with the available genomes at public databases and B) Additional details regarding analysis of the mating type system of *M*. *furfur*.

**Supplementary Figure 1.** IGV screenshots showing the genomic patterns of M. furfur. A) Read mapping of CBS1878 and CBS7019 on the reference genome, showing an example of a region of high genomic variability. B) Read mapping of P1 strains to CBS7982 mitochondrial genome assembly. C) Read mapping of H1 strains to MAT a2 loci of P2, showing the occurrence of a duplication.

**Supplementary Figure 2.** A) Karyotype variation (PFGE) and B) FACS analysis present variation between both hybrid lineages H1 and H2 at the chromosomal and ploidy levels. PFGE gels were run under different conditions (see supplementary materials and methods and supplementary Table 2).

**Supplementary Figure 3.** A) Phylogenetic trees for five nuclear DNA loci. Phylogenetic trees based on the Maximum Likelihood method and Tamura-Nei model with 500 bootstrap replications for A1) internal transcribed spacer (ITS) and A2) the intergenic spacer 1 region (IGS1) of the ribosomal DNA. The IGS1 tree illustrates the suitability of this locus for discrimination between the four studied lineages. The ITS tree showcases the lack of discrimination between both hybrid lineages and the P2 lineage. Additional phylogenetic trees are presented for protein coding genes A3) β-tubulin, A4) TEF1 and A5) CHS2, illustrating the potential difficulties when sequencing protein coding genes for hybrid strains as at multiple spots in the chromatograms, two different base peaks are present, representing both parental haplotypes. Therefore, the sequences of the protein coding were phased into two copies, based on their parental haplotypes, and B) Dendrogram based on MALDI-TOF MS generated Main Spectra (MSPs) as a measure of general phenotypic variation at the proteomic level, illustrating variation between the various lineages.

**Supplementary Figure 4.** Coverage plot obtained from sppIDer pipeline (78) for the H1 strains (left) and H2 strains (right) used for whole-genome sequencing data analysis when aligned to a combined reference genome of CBS9595 (P1 lineage, red) and CBS14141 (P2 lineage, blue). Dashed lines divide the different chromosomes.

**Supplementary Materials and Methods**. Descriptions of Pulsed-Field Gel Electrophoresis (PFGE), Fluorescence-activated cell sorting (FACS), Matrix-Assisted Laser Desorption Ionization-time Of Flight Mass Spectrometry (MALDI-TOF MS), Microscopy (Light microscopy – cell size measurements), Physiology, and Sanger sequencing and phylogenetic analysis.

## Notes

### Competing Interest Statement

The authors have declared no competing interest.

### Summary of Updates

This second revision has a few small structural changes to improve readability and includes some reorganizing of supplementary files.

