## Supplementary File 1 for "Multiple hybridization events punctuate the evolutionary trajectory of *Malassezia furfur*"

**Supplementary File 1A.** **Genome assembly of *M. furfur* parental lineages and comparison with the genomes available in public databases.**

A major goal of this project was to explore the genome of *M. furfur* hybrids and determine their evolutionary path. Given the presence of isolates corresponding to their putative parental lineages among the isolates (see Figure 1 of the manuscript), the best approach to analyze these genomes was by read mapping of their Illumina reads against a reference genome comprising the two parental subgenomes. At the beginning of this project, the only genomes available for the putative parental lineages were those of CBS7982 and CBC14141 (accession numbers: [GCA_001265065.1](https://www.ncbi.nlm.nih.gov/assembly/GCA_001265065.1) and [GCA_001265045.1](https://www.ncbi.nlm.nih.gov/assembly/GCA_001265045.1) (Wu et al. 2015)), corresponding to approximately 7.8 Mb distributed in 1,694 and 2,092 scaffolds, respectively. In order to obtain more contiguous assemblies for these two lineages, we sequenced these genomes with long-read sequencing technologies, as well as the genome of an additional strain of P1 (CBS9595). DNA sequencing and assembly protocols are described in detail in the Materials and Methods section of the manuscript.

**Genome assembly of *M. furfur* parental lineages**

A *k*-mer based analysis (see Materials and Methods for details) pointed to an estimated genome size of 7.26 Mb for CBS7982, 7.83 Mb for CBS9595, and 8.3 Mb for CBS14141 (Table 2 of the manuscript). The genome assembly process was performed independently following various strategies in parallel combining both short- and long-read sequencing data for each of the three strains (see Materials and Methods for more details). In the end, the best genome assembly for each strain was selected based on the N50, assembly fragmentation, and estimated completeness. In the case of CBS7982, the final assembly comprised 8.1 Mb divided in 14 scaffolds, with an N50 of 1 621 469 bp, corresponding to an estimated assembly completeness of 99.9%, thus corresponding to a significant improvement over the previously available assembly. Regarding the genome of the other strain of the parental lineage P1, CBS9595, the final assembly comprised 8.1 Mb divided in 8 scaffolds, with an N50 of 1,622,862 bp, corresponding to an estimated assembly completeness of 99.9%. Given the robust assembly features of both strains, but the higher contiguity observed in CBS9595, this last assembly was considered the best candidate for the reference of P1 lineage. Before finalizing this decision, it was essential to guarantee that it was a good representative of the lineage. Therefore, we did a *k*-mer based comparison between the available libraries for the remaining strains of the P1 lineage and CBS9595 assembly (see Materials and Methods for more details). This analysis revealed that this assembly represented more than 97% of the *k-*mers of all the P1 strains, hence it was selected as the representative of this lineage for further analysis.

For the putative parental lineage P2, we assembled the genome of CBS14141, following various strategies in parallel combining both short- and long-read sequencing data (see Materials and Methods for more details). The final assembly comprised 8.2 Mb divided in 9 scaffolds, with an N50 of 1 642 932 bp, corresponding to an estimated assembly completeness of 99.8%. This is a significant improvement over the initially available assembly. Of note, only 77.2% of CBS14141 Illumina reads aligned to the assembly, which is in high contrast with the estimated assembly completeness. In previous studies from our group on the analysis of the genome of *Diutina* *(Candida) rugosa*, we observed a similar pattern (Mixão et al. 2019). In this case, only 60% of the reads aligned in the assembly of the nuclear genome, because the remaining 40% corresponded to the mitochondrial genome. Therefore, we hypothesized that a similar situation could be occurring with CBS14141. Hence, we mapped the Illumina reads of this sample simultaneously to the nuclear genome assembly that we generated and the mitochondrial genome assembly that was available in NCBI (accession number: KY911085.1). This increased the number of mapped reads to 99.4%, indicating that the Illumina library indeed includes a high proportion of mitochondrial content, and therefore the low percentage of mapping reads is not an indicator of a low assembly quality. Similar to what was proposed for *D. rugosa*, we are not certain if this observation is related to a high mitochondrial content in this sample, or to a bias introduced during library preparation. However, it is noteworthy that *M. furfur* CBS14141 (retrieved from NCBI) and *D. rugosa* libraries were prepared and sequenced in different laboratories. Similarly to what we did for P1 lineage, we performed a *k-*mer based comparison between the libraries of P2 lineage and the final CBS14141 genome assembly to assess whether it is a good representative of these strains. This analysis revealed that this genome assembly represented more than 97% of the *k-*mers of all the P2 strains, hence it could serve as a representative of the lineage.

**Annotation of *M. furfur* P1 and P2 assemblies**

We next performed genome annotation for the two genome assemblies that would be used in further analysis (CBS9595 and CBS14141). These results also served as an additional assembly quality control measure. As noted in the Materials and Methods section of the manuscript, annotation was performed with Augustus Web-server (Hoff and Stanke 2019; Stanke and Morgenstern 2005). As Augustus default training sets do not include a *Malassezia* species, we decided that the best approach would be to use a proteome from the NCBI database to train Augustus. Therefore, we decided to download the proteome of *Malassezia restricta* (accession number: ASM329048v1)*, Malassezia sympodialis* (accession number: ASM34930v2) and *Malassezia globosa* (accession number: ASM18169v1), and assess their completeness with BUSCO v4 using the Basidiomycota database (Seppey, Manni, and Zdobnov 2019). The estimated completeness was 95.8%, 73.7% and 80.2%, respectively. Considering these results, we decided to train Augustus with the proteome of *M. restricta*. Afterwards, we used this training set to annotate the genome assembly of CBS9595 and CBS14141. We predicted 4 376 and 4 441 protein-coding genes, respectively. This corresponds to an estimated proteome completeness of 96.1% and 95.4% for CBS9595 and CBS14141, respectively, which is also a good indicator of the assembly quality.

**Supplementary File 1B. *M. furfur* possesses the genetic machinery of a pseudo-bipolar mating system**

To understand the origin of the hybridization events that formed the H1 and H2 lineages, the previously described *Malassezia* mating-type genes (Xu *et al.* 2007; Gioti *et al.* 2013; Wu *et al.* 2015). were investigated here for *M. furfur*. The *MAT* genes of *M. furfur* haploid strains were searched with tBLASTn using as query the previously reported *MAT* genes of two opposite mating-types of *M. sympodialis* (strain CBS12506, *MAT a1b1*; strain ATCC44340, *MAT a2b2*). Two *MAT* *a* loci were identified in the haploid *M. furfur* strains analyzed belonging to P1 and P2 lineages, and they were designated as *a1* and *a2* according to the closest *M. sympodialis* orthologs (Supplementary Table 6). Both the Mfa1 and Pra1 proteins of the P/R locus of *M. sympodialis* CBS12506 found significant matches in the PacBio assemblies of *M. furfur* strains CBS14139 (syn. JPLK13), CBS8735, and PM315, which were designated as *a1*. For strains CBS14141 and CBS7982, tBLASTn analysis revealed higher similarity with the pheromone receptor of *M. sympodialis* ATCC44340, and thus are predicted to be *MAT* *a2*, but orthologs of the pheromone were not found in these cases. Therefore, following the *MAT* structure of other *Malassezia* strains (Xu *et al.* 2007; Gioti *et al.* 2013; Triana *et al.* 2015) and using *M. furfur* CBS14141 as the reference, a 1500-bp region from the identified *pra2* gene was manually screened for the sequence predicted to encode the *mfa2* gene, based on two main conserved features: the presence of a cysteine codon three codons prior to the stop codon, and the presence of a CAAX motif in the protein sequence. These features, combined with the potential ATG start codons and stop codons, allowed us to define a region of 153 bp that is predicted to be the *mfa2*-encoding pheromone gene and that completed the *MAT* *a2* locus of *M. furfur*. This prediction was further confirmed by RNA-seq data. For strain CBS9369, high *E*-values were obtained using *M. sympodialis* proteins as a query, and therefore its characterization as *MAT* *a1* was defined after comparison with the predicted *MAT* *a1* and *a2* loci for *M. furfur* strains CBS14139 (syn. JPLK13) and CBS14141, respectively (Supplementary Table 6). The availability of annotated *MAT* genes in *M. sympodialis*, *M. globosa,* and *M. pachydermatis* (Xu *et al.* 2007; Gioti *et al.* 2013; Wu *et al.* 2015), and the availability of RNA-seq for *M. furfur* CBS14141, enabled the identification of all ORFs, with the exception of the *pra1* gene for which transcriptomic data are not as yet available.

Similarly, the *M. furfur bE* and *bW* genes of the *MAT* *b* loci encoding homeodomain transcription factors were initially identified as orthologs of *M. sympodialis* bE and bW, but due to the high similarity, their identity could not be unambiguously assigned based on the BLAST outcome (Supplementary Table 6). Phylogenetic analysis of the concatenated alignments of the predicted proteins revealed the presence of two *MAT* *b* loci defined as *b1* and *b2*, which did not cluster with the *MAT* *b1, b2,* and *b3* alleles of *M. sympodialis* (Gioti *et al.* 2013) (data not shown).

Of note, for *M. furfur* strain CBS9369 the pseudo-bipolar *MAT* structure could not be confirmed. This is likely due to the more fragmented nature of the genome assembly, which resulted in the *MAT a1* and *MAT b2* loci being located on different scaffolds sharing high similarity (*E*-values as 0.0 following BLAST analyses) with the respective *MAT* loci identified in the other *M. furfur* strains.

For the two *MAT* *a locus* genes DNA sequence similarity is 58.74% between the *pra1* and *pra2* genes, and 61.48% between the *mfa1* and *mfa2* genes. The *MAT* *b1* and *b2* loci are highly syntenic, and in this case the *bE* and *bW* genes are also divergently oriented, as in other *Malassezia* species (Xu *et al.* 2007; Gioti *et al.* 2013; Wu *et al.* 2015) (Figure 5). The HD genes of the *MAT* *b* loci share high sequence identity, with 88.42% between the *bE1* and *bE2* genes and 87.17% between the *bW1* and *bW2* genes.
