## Supplementary materials and methods for "Multiple hybridization events punctuate the evolutionary trajectory of *Malassezia furfur*"

**Supplementary Materials & Methods**

**Pulsed-Field Gel Electrophoresis (PFGE)**

PFGE was performed according to (Boekhout *et al.* 1998) with electrophoresis settings according as listed below.

| **Electrophoresis parameter** | **Gel A** | **Gel B** | **Gel C** |
| --- | --- | --- | --- |
| Run time 1 (h) | 10 | 36 | 36 |
| Pulse times (s) | 60 - 60 | 300 – 300 | 300 - 600 |
| Run time 2 (h) | 34 | 36 | 48 |
| Pulse times (s) | 300 - 400 | 300 – 600 | 600 - 900 |
| Initial current (mA) | 70 | 130 |  |
| End current (mA) | 70 | 170 |  |
| Voltage (V) | 120 | 100 | 80 |
| Temperature (C) | 14 | 14 | 14 |
| Gel concentration (%) | 1 | 1 | 1 |
| Bufferconcentration TBE | 0.5x | 0.5xx | 0.5x |

**Fluorescence-activated cell sorting (FACS)**

Ploidy was determined by fluorescence-activated cell sorting (FACS) analysis as described by (Lengeler *et al.* 2001) with minor modifications. *M*. *furfur* strains were grown for 3 days on mDixon agar at 30 °C, cells were collected in water, washed one time with PBS and fixed in 70 % ethanol at 4 °C overnight. To stain nuclei, fixed cells were washed with NS buffer (10 mM Tris-HCl, pH 7.6, 250 mM sucrose, 1 mM EDTA, pH 8.0, 1 mM MgCl2, 0.1 mM CaCl2, 0.1 mM ZnCl2) and stained with propidium iodide (10 mg/ml) in 0.2 ml of NS buffer containing RNase A (1 mg/ml) at 4 °C overnight. Stained cells (50 μl) were added to 0.5 mL of Tris-PI mix [482 µl 1M Tris pH 7.5+18 µl Propidium Iodide (1 µg/µl)]. Flow cytometry was performed on 20,000 cells with slow laser scan, on the FL1 channel with a Becton-Dickinson FACScan.

**Matrix-Assisted Laser Desorption Ionization-time Of Flight Mass Spectrometry (MALDI-TOF MS)**

Strains were cultured on mDixon agar for 48 h at 30 °C after which sample preparation was carried out with the ethanol/formic acid protein extraction method, following the Bruker Daltonics GmbH protocol, using two loops of Malassezia cells (1 µL volume, sterile inoculation loop). Additional steps were executed according to the methods described by Kolecka and colleagues (Kolecka *et al.* 2014). Clustering of Main Mass Mpectra (MSPs) was performed using MBT Compass Exlorer (v 4.1, Bruker Daltonics GmbH, Bremen, Germany) using default parameters.

**Microscopy (Light microscopy – cell size measurements)**

*M*. *furfur* cells were grown on mDA for 72 h at 30 °C and three loops of cells were separately suspended in water and length and width of 50 randomly picked individual cells was assessed under 60 x magnification with a Zeiss Axio microscope (Jena, Germany).

**Physiology**

Physiology experiments were performed according to Guého et al. (Guého-Kellermann *et al.* 2010). Temperature growth experiments were performed, using 2d-old cultures on mDixon making a suspension in 1 ml sterile demi water using a 10 μl sterile disposable inoculation loop. From this suspension, 2 drops of 3 μl each, were applied on the tested culture media and observed for growth at the respective temperatures.

**Sanger sequencing and phylogenetic analysis**

In order to further support the lineage determination and genetic variation as established by AFLP, for the 22 strains in this study, four loci were sequenced using Sanger sequencing: the ITS region of the ribosomal DNA, and parts of the protein coding genes chitin synthase (CHS2), translation elongation factor EF-1α (TEF1), and β-tubulin. PCR-reactions were performed in 25 μL volumes, with 2.5 μL 10x NH4 Reaction Buffer, 1.5 mM MgCl2, 250 μM of each dNTP, 0.3 μM (Meridian Bioscience, Cincinnati, USA) of each primer, 0.5 U of BioTaq (Meridian Bioscience, Cincinnati, USA) and 1 μL of template DNA. PCR conditions were general with the exception of the annealing temperature: 96 °C for 5 min; 35 cycles of 96 °C for 30 s, annealing temperature (Ta) °C for 45 s, and 72 °C for 70 s; and a final extension at 72 °C for 10 min. Primers’ sequences and annealing temperatures (Ta) for the various loci are listed below.

| **Locus** | **Ta (°C)** | **Primer name** | **Primer sequence 5’ > 3’** |
| --- | --- | --- | --- |
| ITS | 55 | ITS5 | GGAAGTAAAAGTCGTAACAAGG |
|  |  | ITS4 | TCCTCCGCTTATTGATATGC |
| TEF1 | 55 | EF1-983F | GCY CCY GGH CAY CGT GAY TTY AT |
|  |  | EF1-1567R | ACH GTR CCR ATA CCA CCR ATC TT |
| CHS2 | 55 | ChiSyn2f | CTG AAG CTT CAN ATG TAY AAY GAR GAY |
|  |  | ChiSyn2r | GTT CTC GAG YTT RTAYTC RAA RTT YTG |
| β-tubulin | 50 | F-β-tub | CARGCYGGTCARTGYGGTAACCA |
|  |  | F- βtub4r | GCCTCAGTRAAYTCCATYTCRTCCAT |

All PCR products were sequenced in both directions with the primers that were also used for the amplification. Raw sequencing data was assessed and consensus sequences created in Seqman Pro^TM^ (v. 9.0.4 (39), DNASTAR, Inc., Madison, USA). Sequences were aligned and trimmed in MEGAX (v 10.1.7) (Kumar *et al.* 2018) using MUSCLE (Edgar 2004) and for protein coding genes, sequences for hybrid strains were manually phased according to their parental copies, with reassessment of chromatograms for polymorphic sites. Finally, phylogenetic trees were created with MEGAX (Kumar *et al.* 2018) as well, using the Maximum Likelihood method and Tamura-Nei model and 500 bootstrap replications.
