## Supplementary figures and images for "Multiple hybridization events punctuate the evolutionary trajectory of *Malassezia furfur*"

### Supplementary Figure 1

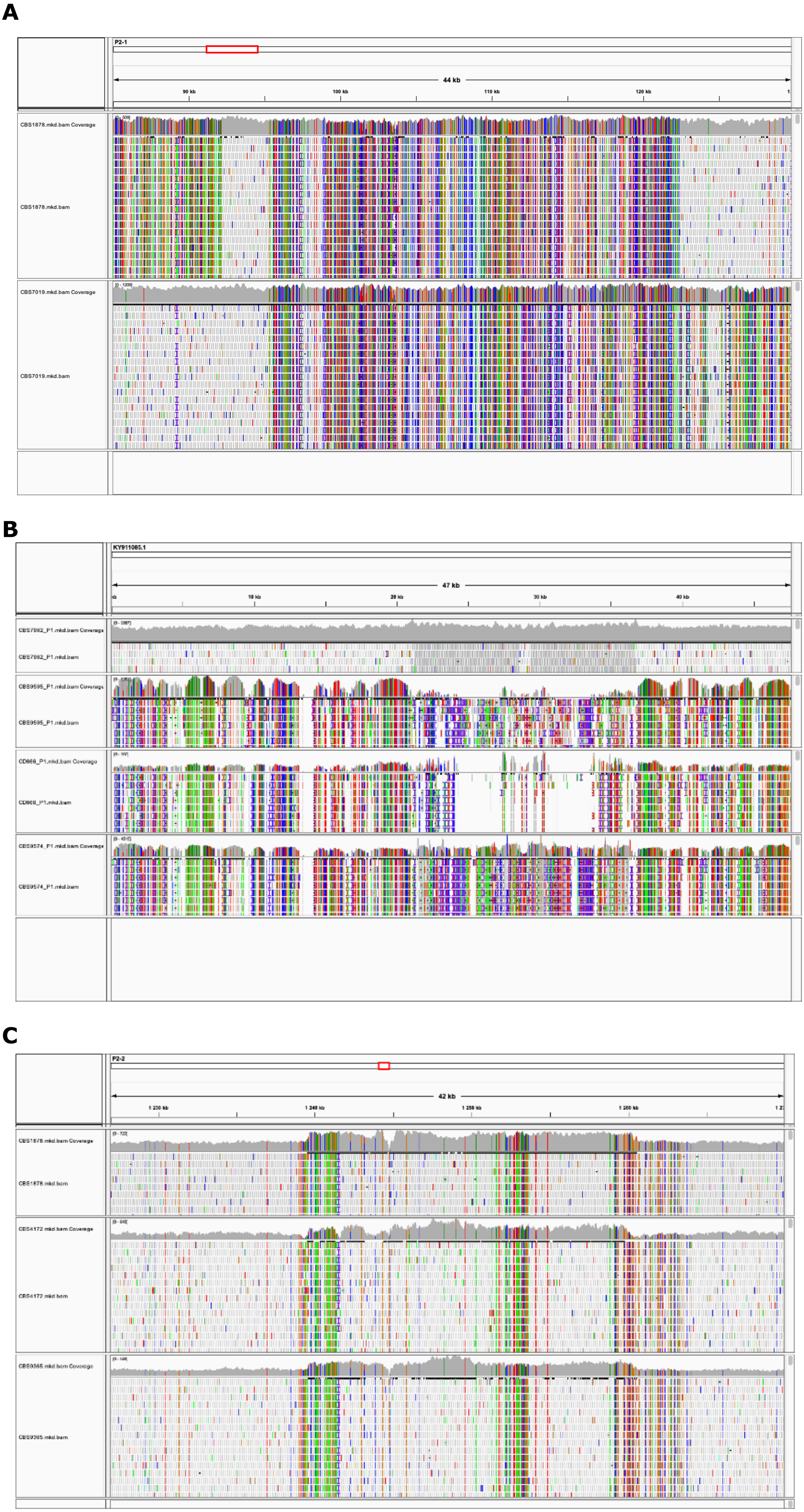

### Supplementary Figure 2

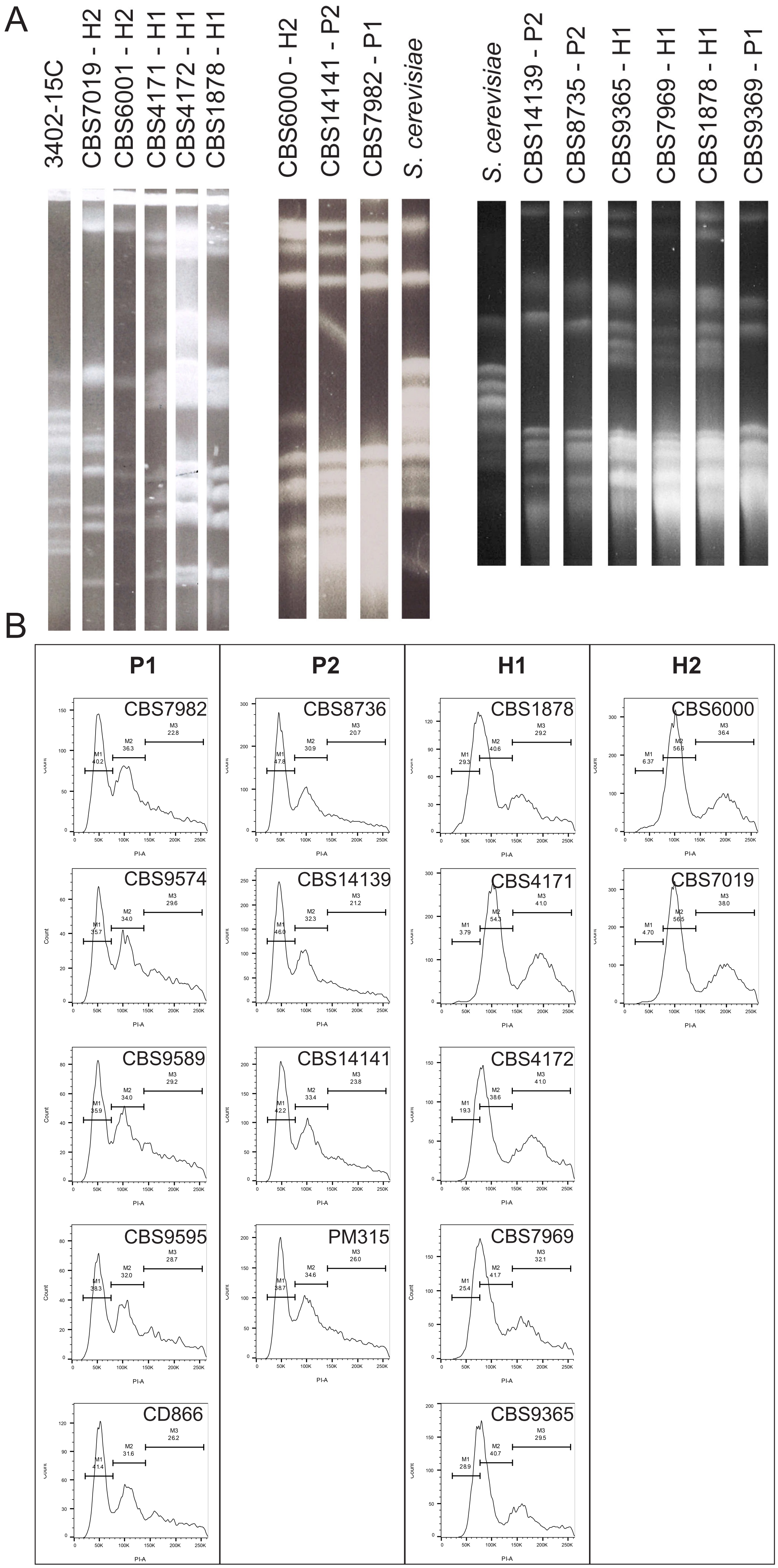

### Supplementary Figure 3

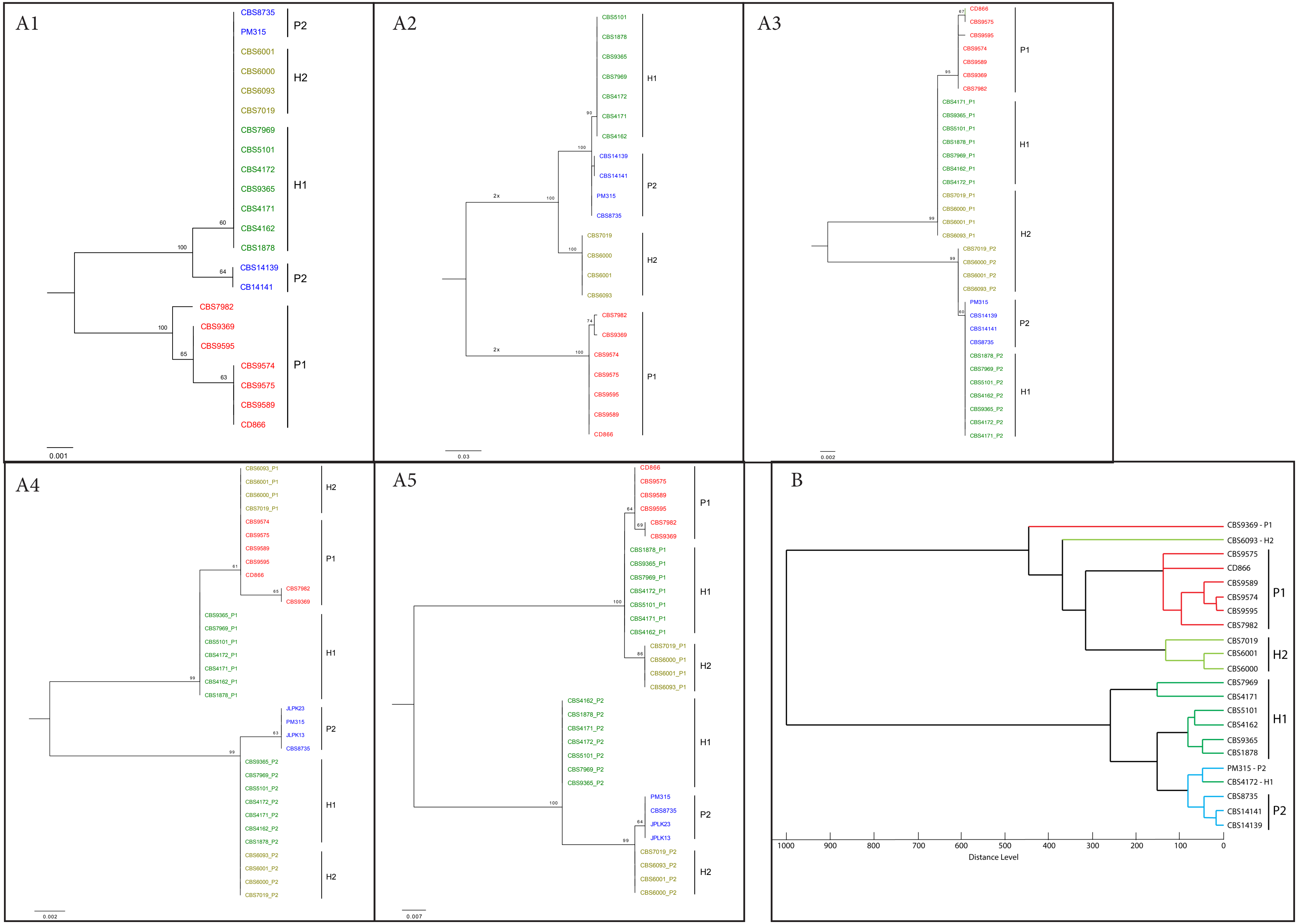

### Supplementary Figure 4

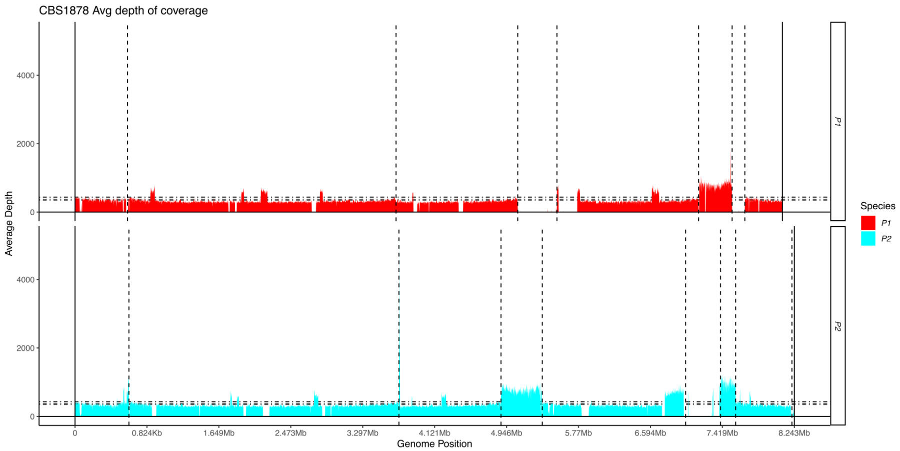

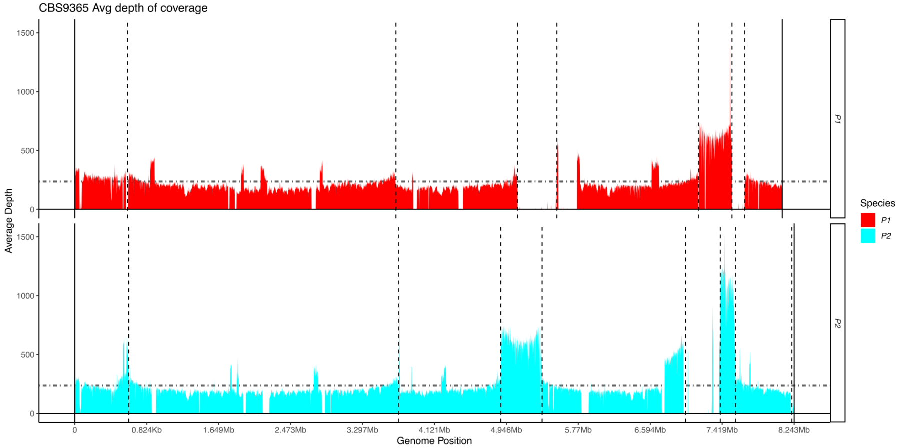

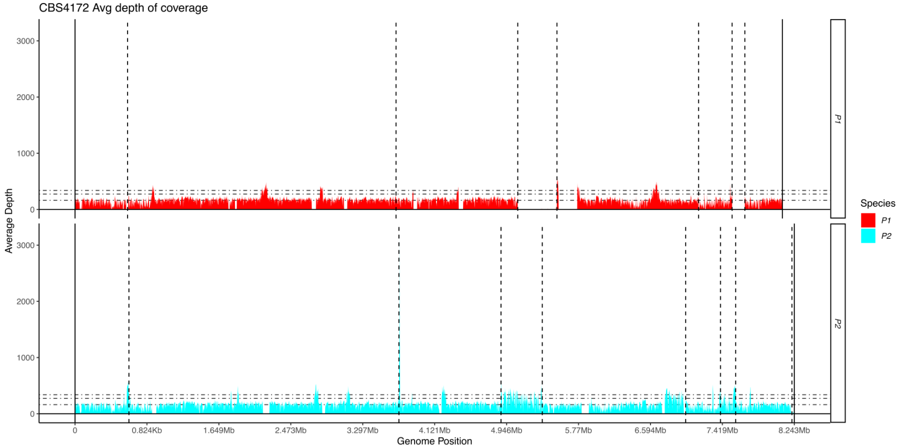

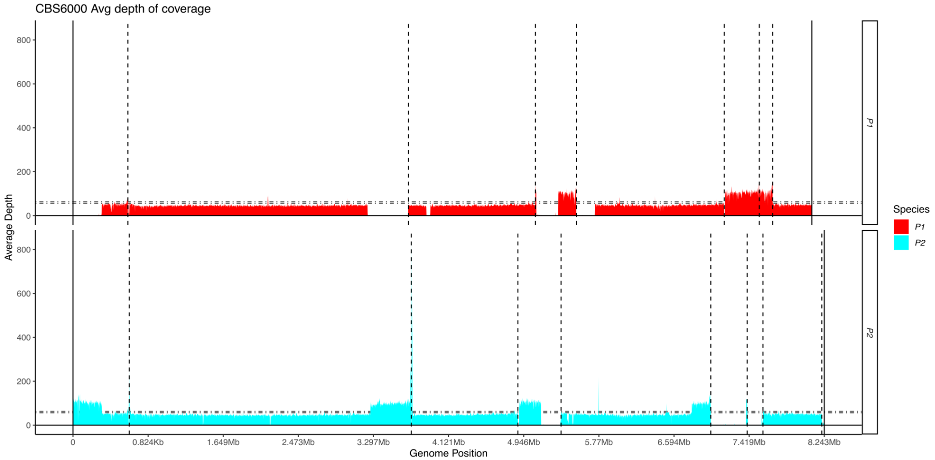

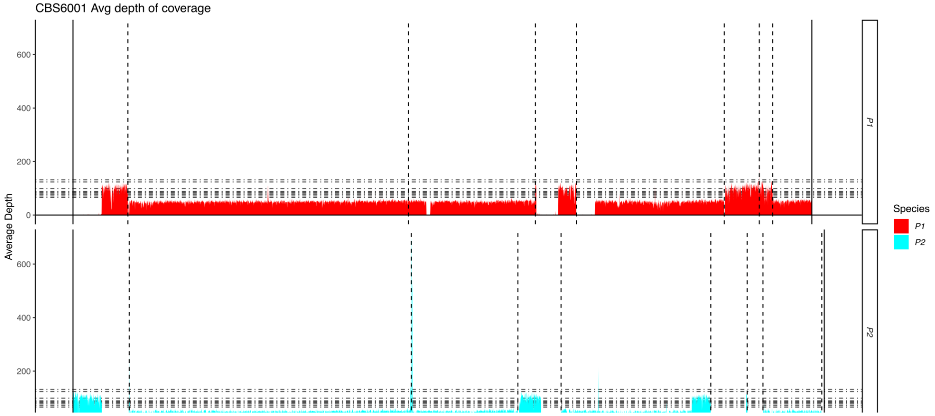

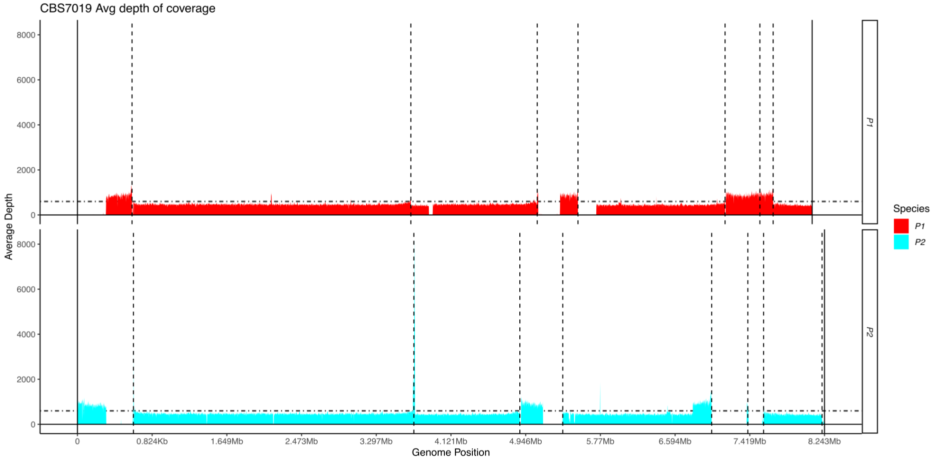
